# Unraveling long-term trends and drivers of fish biodiversity change using environmental DNA metabarcoding of archived samples

**DOI:** 10.64898/2025.12.23.696188

**Authors:** Robin Schütz, Martin Friedrichs-Manthey, Till-Hendrik Macher, Arne J. Beermann, Jens Arle, Jan Koschorreck, Florian Leese

**Affiliations:** University of Duisburg-Essen, Aquatic Ecosystem Research, Universitätsstr. 5, 45141 Essen, Germany; Institute of Biodiversity, Friedrich Schiller University Jena, Dornburger Straße 159, 07743 Jena, Germany; Helmholtz Centre for Environmental Research - UFZ, Department of Ecosystem Services, Permoserstr. 15, 04318 Leipzig, Germany; German Centre for Integrative Biodiversity Research (iDiv) Halle-Jena-Leipzig, Puschstr. 4, 04103 Leipzig, Germany; Trier University, Department of Biogeography, Universitätsring 15, 54296 Trier, Germany; University of Duisburg-Essen, Centre for Water and Environmental Research (ZWU), Universitätsstr. 3, 45141 Essen, Germany; German Environment Agency, Wörlitzer Platz 1, 06844 Dessau-Roßlau, Germany

**Keywords:** eDNA, monitoring, time-series, environmental specimen bank, multiple stressors

## Abstract

Long-term biodiversity data are critical for assessing, understanding, and predicting ecosystem change globally, but are often limited by their availability. We present the longest eDNA metabarcoding time series for freshwater fishes based on archived suspended particulate matter (SPM) from Germany’s Environmental Specimen Bank, covering 17 years (2005-2021) across six major European river systems (Rhine, Danube, Elbe, and their tributaries). From 211 annual samples collected according to highly standardized procedures at 13 sites, we detected 63 fish and lamprey species and traced their spatio-temporal trajectories. eDNA metabarcoding captured significantly declining trends at multiple sites with no recoveries. Common and sensitive species declined at nearly half of sites while invasive species spread. Site occupancy and relative read abundance trends were correlated, providing semi-quantitative indicators. Alpha and beta diversity shifted over time and Bayesian hierarchical models revealed that 19% of species showed significant declines in relative read abundance. The eDNA-derived trends showed strong concordance with long-term regulatory fish monitoring data, validating the patterns across methodological approaches. Random forest models and multivariate analyses identified multiple anthropogenic pollutants, measured from the same SPM samples, as drivers of biodiversity patterns. Pesticides, heavy metals, temperature, and fine sediment negatively affected species richness, with site-specific stressor combinations explaining 29% of community variation. Our results demonstrate that archived SPM, originally collected for chemical monitoring, provides a reliable, dual-purpose monitoring matrix integrating pollution and biodiversity trend assessments simultaneously. This approach enables holistic monitoring of global freshwater ecosystem change and can provide an early warning framework supporting evidence-based environmental management.

## Introduction

Time series data plays a crucial role in global biodiversity research and monitoring, providing detailed insights into the long-term dynamics of ecosystems. Unlike individual surveys, systematic long-term monitoring enables the detection of gradual shifts, rapid regime changes, long term-trends and ecological transitions driven by factors such as climate change, habitat alteration, pollution, and various other anthropogenic stressors (Dornelas et al., 2013). Such continuous data collection is particularly important in freshwater ecosystems, which are subject to rapid and complex environmental changes (Kaijser et al., 2025; Tickner et al., 2020). For example, long-term data have shown that initial improvements in water quality and biodiversity recovery within European river systems have come to a halt in recent years due to persistent anthropogenic pressures (Haase et al., 2023).

One key group of interest in these aquatic ecosystems are fish, which serve as sensitive bioindicators, play central roles in trophic interactions, and are of economic relevance. Due to their often complex life histories and narrow habitat requirements, even slight alterations in water quality, flow regimes, or habitat structure can lead to significant shifts in fish community composition and behavior (Radinger et al., 2019). A recent global meta-analysis quantified stressor-response relationships for riverine organisms, showing that fish communities typically respond negatively to salinity, oxygen depletion, sediment accumulation, and flow cessation, while responses to nutrient enrichment and warming are more variable (Kaijser et al., 2025). Understanding these biological responses can provide an early warning system of ecosystem health that is not always captured by chemical or physical assessments alone. Consequently, freshwater fish are included as protection goals and indicators in regulatory monitoring programs such as the European Union Water Framework Directive (Birk et al., 2012).

Despite the urgent need for comprehensive surveys, the availability of large-scale standardized spatio-temporal fish monitoring data is limited by several challenges, including low temporal resolution, limited spatial coverage, high costs, labor intensiveness, and a lack of standardized protocols (Haase et al., 2018). In this context, environmental DNA (eDNA) metabarcoding has emerged as a promising tool. eDNA-based methods involve collecting genetic material released by organisms into their environment (e.g., water, or sediment), enabling rapid, non-invasive, and cost-effective detection of species across the tree of life and including cryptic, rare, and invasive species (Beng & Corlett, 2020; Macher, Schütz, et al., 2021). These methods not only enable a more holistic monitoring approach but also allow for increased sampling frequency and broader geographic coverage as it can be employed at lower cost per sample (Seymour et al., 2021). Recent studies have demonstrated that integrating aquatic eDNA approaches into monitoring frameworks significantly enhances the resolution and comparability of time-series datasets. For instance, annual eDNA analyses have captured seasonal and interannual fluctuations in lake assemblages (Bista et al., 2017), while long-term marine studies have revealed nuanced community trends (Djurhuus et al., 2020). Similarly, riverine eDNA investigations have detected dynamic shifts in multi-trophic communities in response to eco-hydrological variation (Liang et al., 2022). These findings highlight eDNA’s potential to streamline data collection and support robust assessments across diverse aquatic environments (Altermatt et al., 2025). Integrating fine-scaled time series data using scalable methods like eDNA metabarcoding also offers a later historical perspective on the obtained data and allows for predictive insights essential for safeguarding ecosystem resilience and targeting restoration and conservation efforts (Carraro et al., 2020; Schütz et al., 2025; Sullivan et al., 2025).

In densely populated regions like Central Europe, monitoring freshwater fish is especially critical. The intense pressures from urbanization, intensive agriculture, and industrial activities lead to habitat degradation, fragmented river connectivity, and declining water quality, all of which have a pronounced impact on aquatic ecosystem health (Hupalo et al., 2022; Schweizer et al., 2022). In Germany, the status of freshwater fish and lampreys is of growing concern as many native species are now listed as endangered or threatened in the national red list (Freyhof et al., 2023), and migratory species, such as the European river lamprey (*Lampetra fluviatilis*), are vulnerable due to discontinued river corridors caused by infrastructural barriers like dams (Lothian et al., 2024). Moreover, invasive goby species, such as the round goby (*Neogobius melanostomus*), bighead goby (*Ponticola kessleri*), and western tubenose goby (*Proterorhinus semilunaris*), have become established in Central European waterways in the last two decades. Studies in the Neckar River demonstrated that the spread of these Ponto-Caspian gobies is linked to the decline in native fish populations, including species like the stone loach (*Barbatula barbatula*) and gudgeon (*Gobio gobio*), thereby altering community dynamics (Gaye-Siessegger et al., 2022). Research on the functional ecology of these invasive gobies further underscores their disruptive effects on trophic interactions and overall ecosystem processes (Grabowska et al., 2023). Here, long-term and fine scaled monitoring campaigns are key to understanding species distribution, assessing their impact, and ensuring effective management. A recent study has investigated fish abundance data from regulatory monitoring from 2004 to 2020 in rivers in Central and Northern Germany (Friedrichs-Manthey et al., 2024). Their work revealed that while several native fish species exhibited significant declines in abundance, several non-native species showed increasing trends, pointing to progressively altered fish communities under the influence of anthropogenic drivers.

As eDNA metabarcoding was first introduced for biodiversity assessment in 2012 (Thomsen et al., 2012), older eDNA data are scarcely available. Provided that samples are collected and stored in a quality-assuring manner, archived environmental samples can be a source to obtain time series for retrospective biodiversity analysis (Krehenwinkel et al., 2022; Zizka et al., 2022). The German Environmental Specimen Bank (ESB) is a long-term infrastructure for human and ecosystem health monitoring established in the 1980s that has been collecting and storing different environmental matrices for decades in support of German environment policy and regulation. These samples, so far mainly used for pollution trend monitoring, can now be analyzed with genetic methods as they are stored at - 150°C above liquid nitrogen (Junk et al., 2025; Zizka et al., 2022). Among the archived sample matrices are suspended particulate matter (SPM) from national long-term sampling sites in Germany’s largest waterways. The oldest samples originate from 2005, when SPM sampling started, but eDNA metabarcoding was not yet established. The first pilot study successfully amplified and sequenced fish DNA from these archived samples (Díaz et al., 2020). Building on that, further tests were carried out optimizing the extraction and replication strategy so that the archived SPM material could be used effectively for the detection of fish and invertebrates (Schütz et al., 2025).

This study explores retrospective eDNA metabarcoding from archived SPM samples from 2005-2021 to assess changes in fish communities from 6 major European rivers and identify environmental drivers. For this, we use both presence-absence as well as relative read abundance data. Specifically, we address three hypotheses:

1. The eDNA SPM time series data (2005-2021) captures broader spatial and temporal trends in fish alpha and beta diversity from community trends to detailed patterns of key groups, as well as species-specific trends.
2. Spatiotemporal patterns of fish biodiversity change identified with the eDNA metabarcoding dataset generally align with those observed in regulatory fish monitoring.
3. Physiochemical environmental parameters obtained majorly from the same SPM samples explain a significant variation in fish species richness and community composition, with distinct effects of pollutants, nutrients, and physical gradients.

By testing these hypotheses, we will evaluate the general potential of eDNA metabarcoding of archived SPM to support biodiversity trend monitoring for fish species in rivers, identify sites and species of concern, identify key environmental stressors, and determine whether archived SPM metabarcoding can provide supportive management-relevant insights beyond those generated through conventional data collection.

## Methods

### Sampling

The SPM samples were retrieved from the cryo-archive of the German Environmental Specimen Bank (ESB) which is operated on behalf of the German Environment Agency (Umweltbundesamt) in support of environmental policy and regulation. Sampling, sample processing and archiving follow the ESB standardized procedures (Schulze et al., 2007). At the sampling sites, SPM accumulates constantly in stainless steel sedimentation boxes which are emptied monthly, stirred and sieved to <2 mm particles size, and freeze dried on site. The 12 monthly samples of one year are then stored as a single pooled sample in cryogenic tanks above liquid nitrogen (-150 °C). The ultra-low temperatures in the storage tanks below the glass transition temperature of water and an inert-gas atmosphere prevent oxidation processes ensuring that chemical processes in the samples are reduced to a minimum. In total, we used 211 annual homogenates from 13 long-term sampling sites originating from six rivers, including the largest waterways and catchments of Central Europe (Rhine, Danube and Elbe with their tributaries Mulde, Saale and Saar). The SPM time series covers 17 years (2005-2021), except for the rivers Danube (two sites, 2009-2021) and Saar (two sites, 2006-2021) where SPM sampling was established later.

### DNA extraction, PCR and sequencing

The processing of the samples was performed in a dedicated eDNA laboratory (UV lights, sterile benches, coveralls, gloves, and face masks). To maximize species detection, each annual sample was extracted in duplicates following a previously validated extraction method for SPM based on GuHCI, silica membrane columns, and a magnetic bead clean-up (Buchner, 2022; Schütz et al., 2025). Each extraction duplicate was subsequently amplified with four PCR replicates, using a two-step PCR protocol. The PCR workflow and conditions followed Schütz et al., 2025. In the first step, tagged versions of the tele02 primer (Taberlet et al., 2018) were used to amplify a fragment of 12S gene, as these primers are currently the most suitable for European freshwater fish (Macher et al., 2023). In the second PCR step, Illumina sequence adapters with a dual twin-indexing system were added (Bohmann et al., 2022). Subsequently, normalized PCR products were pooled into one library, evaluated on a Fragment Analyzer (High Sensitivity NGS Fragment Analysis Kit, Advanced Analytical, Ankeny, USA) and sent for sequencing on an Illumina NovaSeq 6000 (paired-end 2 × 150 bp) at GENEWIZ, Leipzig, Germany.

### Raw data analysis

Raw reads were received as demultiplexed fastq files. All samples were processed with the APSCALE-GUI pipeline v1.1.6 (Buchner et al., 2022). Initially, raw reads were paired-end merged using default settings (maxdiffpct□=□25, maxdiffs□=□199, minovlen□=□5). Then the primer sequences were trimmed from reads. Subsequently, reads were quality filtered (maxEE□=□1) and only reads with a length of 10 bp below and above the target fragment length (tele02 = 167 bp) were retained. Next, OTUs (operational taxonomic units) were clustered with a 97% percentage similarity threshold following the recommendations for this primer (Macher et al.,2023). Taxonomic assignments were performed with the APSCALE-blast v1.1.6 (task: blastn) against the Midori2 v259 srRNA reference databases (Leray et al., 2022). Subsequently, taxonomic assignments were filtered according to similarity thresholds (species ≥□97%, genus ≥□95%, family ≥□90%, order ≥□85%) following the workflow established for the tele02 primer (see Macher et al., 2023). The resulting taxonomy table and the read table were then converted to the TaxonTable format for downstream analyses in TaxonTableTools v1.4.7 (Macher et al., 2021) (Supplementary table 1). First, replicates were merged and only OTUs with reads in at least two of the four replicates were kept. To account for potential contamination, the maximum number of reads observed across all negative controls of each OTU was subtracted from the number of reads for the respective OTU in each sample. Afterwards, the table was filtered for fish (i.e., classes Actinopteri and Hyperoartia). The plausibility of each taxonomic assignment was verified manually and adjusted if, for example, a hybrid (e.g., *Carassius* sp.), multiple assignments (e.g., *Anguilla anguilla*/*rostrata*), or erroneous entry (e.g., *Cottus asper*) was conflicting a species assignment. All marine food fish or pet fish species were discarded, keeping only freshwater and brackish water species that are reported in the wild in Germany (Supplementary table 2).

### Species richness and group trends

To visualize sampling sites and river networks, a map was generated that also included the species richness trends (analyzed using GAMs) and trend indication for different groups (common native, red list, non-native, tolerant, sensitive species, and migratory species) using R Studio (RStudio 2025.05.1+513) and trait information from freshwaterecology.info (Schmidt-Kloiber & Hering, 2019). Overall temporal richness trends and temporal trends of these groups per site were analyzed using Mann-Kendall test for detecting monotonic trends, and GAMs for non-linear trends. The tests were applied on species richness (absolute count of species), proportional species richness (group richness divided by total site richness in a year), and proportional read abundance (group reads divided by total site reads per year) (see Supplement 1.1 for details).

### Correlation between relative reads and site occupancy

To assess the relationship between temporal trends in spatial occurrence and read abundance trends for each species, we calculated site occupancy and relative read abundance of each species using R Studio. Species-specific temporal trends were assessed using Mann-Kendall’s tau. The species were classified into four categories based on their trend significance. Significant trends in occupancy only, in relative read abundance only, in both metrics, or no significant trends. The correlation between occupancy trends and relative abundance trends was assessed using Spearman’s rank correlation (see Supplement 1.3 for details).

### Diversity analysis

Change in community composition over time and space was analyzed in R Studio using both quantitative (relative read abundance) and qualitative (presence/absence) approaches with Bray-Curtis dissimilarity and Jaccard dissimilarity, respectively. Whitin-site temporal turnover was quantified by calculating the average pairwise dissimilarities between samples from the same site across time lags. Spatial heterogeneity was assessed by calculating mean between-site dissimilarity within each year. Temporal trends in dissimilarity metrics were assessed using Kendall’s rank correlation. Further, the beta diversity was partitioned into turnover (species replacement) and nestedness (species loss/gain), both for within-site temporal changes and between-site spatial patterns.

The community diversity was further quantified using Hill numbers which provide a unified framework for measuring species diversity (species richness, Shannon, and Simpson diversity). Here, q=0 allows for mere presence/absence assessment, while q=1 and q=2 incorporate relative abundances focusing on medium abundant (q=1) and more abundant (q=2) signals. The diversity stability was assessed for each site using the coefficient of variation (CV) of Hill number q = 1 across all years. Temporal trends in Hill numbers at each site were evaluated using Kendall’s rank correlation. The species-level temporal trends in relative abundance were analyzed across all sites using Kendall’s correlation (see Supplement 1.2 for details).

### eDNA trend estimates

Temporal trends in fish species relative abundance were analyzed using Bayesian hierarchical models in R Studio, similar to the analysis of Friedrichs-Manthey et al., 2024 with dataset-specific differences such as using relative reads instead of actual count data. For each species with observations from at least two different years, we fitted a mixed-effects model with site as a random intercept. Models used Gaussian error distribution on logit-transformed responses and were fitted using Markov Chain Monte Carlo (MCMC) sampling. Convergence was assessed using R-hat ≤ 1.1. Species trends were classified based on their 95% credible intervals of the year coefficient. Significantly increasing (CI entirely > 0), significantly decreasing (CI entirely < 0), or stable (CI overlaps 0) (see Supplement 1.4 for details).

### Comparison to long-term trend data from regulatory monitoring

To validate our eDNA-derived trends, we compared species-level temporal patterns with those reported from regulatory monitoring across Central and Northern Germany (Friedrichs-Manthey et al., 2024). This comprehensive dataset comprises 5,497 sites sampled at least twice between 2004 and 2020. While the datasets differ in sampling sites, sampling methodology (eDNA vs. catch data), and abundance metrics (relative reads vs. specimen counts), both provide estimates of long-term population trends across generally overlapping geographic regions and time periods. The species names were harmonized between datasets to account for taxonomic inconsistencies, with manual mapping applied where eDNA resolution was limited to genus level or ambiguous assignments. Only species present in both datasets were analyzed. The trend estimates from both methods were expressed as the annual rates of change on the logit scale (see eDNA trend estimate section). Agreement between trajectories was assessed using multiple complementary approaches in R Studio. The directional concordance was given in percent, Cohen’s kappa to quantify directional agreement beyond chance, Spearman’s rank correlation between trend magnitudes, Wilcoxon rank test for systematic bias, and Bland-Altman analysis to assess mean bias and 95% limits of agreement (see Supplement 1.5 for details).

### Physicochemical correlations

Chemistry data were obtained from the same SPM samples used for metabarcoding (N=66) and complemented with additional water parameters (N=17) collected from the same long-term monitoring sites used for routine monitoring (Supplement 1.6). The data is collected routinely by the Environmental Specimen Bank (SPM) and the regulatory river monitoring programs (water) and publicly available (see Supplementary material S1.6.). In total, 83 environmental parameters were analyzed, including multiple known physical and chemical anthropogenic stressors. Annual mean values were calculated for each site and year to match the spatio-temporal resolution of the eDNA data (Supplementary table 3). First, the relationship between species richness and individual environmental parameters was assessed using Spearman’s rank correlation in R Studio. Species-level relationships between relative read abundance and environmental parameters were assessed similarly. Next, multivariate relationships between community composition and chemical gradients were evaluated using Redundancy Analysis (RDA) with Hellinger-transformation for zero handling. Multicollinearity among chemical predictors was assessed using Variance Inflation Factors (VIF). The significance of the overall ordination model and individual axes was assessed using a PERMANOVA (n=999). To distinguish chemical effects from spatial and temporal confounding, we performed partial RDA (pRDA) using site and year as covariates. To evaluate the predictive power of chemical parameters on a site’s species richness, Random Forest (RF) models were fitted. Statistical significance of RF models was assessed using permutation tests (n=999). Observed R² values were compared to null distributions to calculate p-values (see Supplement 1.6 for details).

## Results

### Overall diversity

After bioinformatic processing of raw reads, a total of 966,255,650 reads clustered into 8,008 OTUs. By only keeping OTUs that were present in at least 2 of 4 replicates and subtracting the maximum reads of each OTU present in the negative controls, a total of 847,400,738 reads and 724 OTUs remained. The negative controls had a total of 5,975,168 reads (0.6% of all reads) which were mainly assigned to humans (2,620,956 reads). The final table for all analyses, after discarding OTUs that had less than 85% similarity to the closest reference sequence and filtering only for freshwater fish, contained 205,054,790 reads represented by 310 OTUs. These were assigned to 63 distinct taxa, including OTUs showing a 100% match to two or more plausible species (e.g., *Alosa* sp., *Coregonus* sp., *Lampetra* sp., *Carassius* sp.). The actual species’ richness could therefore be higher. For simplicity, we hereafter refer to all these distinct taxa as “species”. Species richness ranged from 5 to 35 species per site-year (mean ± SD: 18.7 ± 6.2). In general, the detected species (after removing e.g., marine food fish or pet fish) are all reported from Germany according to the German Red List and regulatory monitoring programs conducted by state authorities. Further, the detected species match the known distribution of species in the rivers with e.g., endemic Danube species only being reported there (e.g., *Gymnocephalus schraetser* or *Rutilus pigus*) or estuary species only detected in the river mouth (e.g., *Osmerus eperlanus* or *Pomatoschistus microps*).

### Species richness trends per site

The linear trend analysis identified declining trends at three sites (Blankenese, Weil, Jochenstein), one in each catchment (Elbe, Rhine, Danube). A fourth site (Prossen, Elbe) was marginally non-significant (p=0.052). Using GAMs, significant negative trends were confirmed for Weil, Prossen and Jochenstein, yet more complex patterns were derived showing linear to non-linear trends (Table 1, Figure 1) with the highest Estimated Degree of Freedom (edf) for Weil. The GAM analysis further revealed varying deviance explained ranging from no explanation (<0.001%) to high explanation (64.6%).

**Figure 1:**
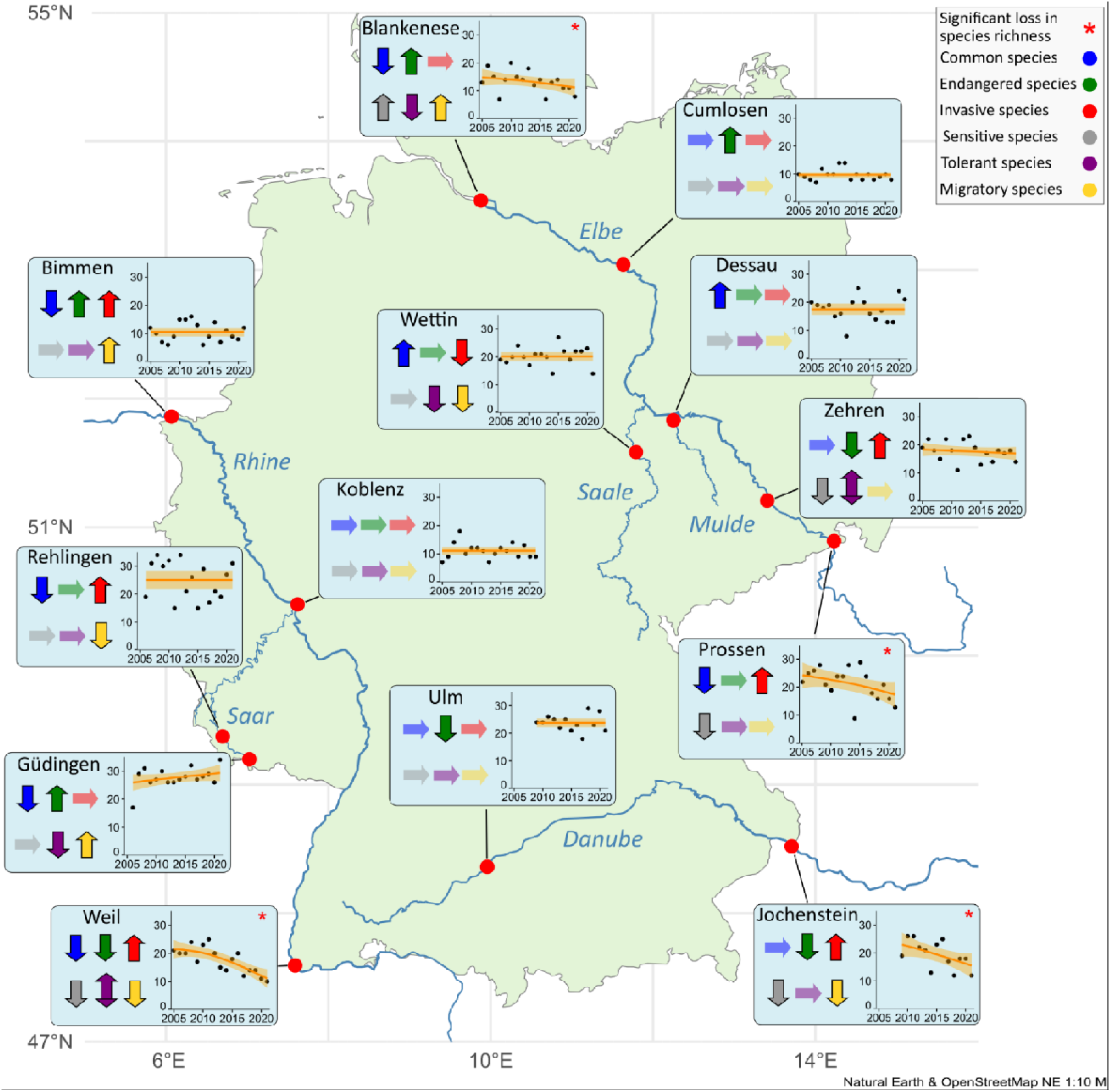
Location of the sampling sites in Germany with Generative Additive Models (GAM) representing the change of species richness over time per site with significance (Mann-Kendall) indicated by an asterisk. Arrows per site indicate a significant trend for the respective group (common species, endangered species, invasive species, sensitive species, tolerant species and migratory species). The upward arrow indicates a significant increase (Mann-Kendall) in at least one of the categories: species richness of the group over time, relative proportion of the groups’ species, and relative read proportion of the group. A downward arrow indicates a significant decrease in one of the categories while a horizontal arrow indicates no significant change over time.

**Tab. 1:** Statistical linear and non-linear analysis of species richness trends per site. Green highlighted results indicate significance, orange indicates almost significant trends.

| River | Site | Mann-Kendall |  | Generalized Additive Model |  |  |
| --- | --- | --- | --- | --- | --- | --- |
|  |  | tau_mk | p_mk | edf | p_gam | dev_expl |
| Elbe | Prossen | -0.361 | 0.052 | 1.124 | 0.016 | 24.662 |
|  | Zehren | -0.262 | 0.168 | 0.102 | 0.294 | 1.942 |
|  | Cumlosen | -0.090 | 0.668 | <0.001 | 0.773 | <0.001 |
|  | Blankenese | -0.382 | 0.041 | 0.844 | 0.075 | 17.973 |
| Mulde | Dessau | -0.075 | 0.709 | <0.001 | 0.886 | <0.001 |
| Saale | Wettin | 0.175 | 0.360 | <0.001 | 0.941 | <0.001 |
| Rhine | Weil | -0.616 | <0.001 | 1.466 | <0.001 | 64.593 |
|  | Koblenz | -0.054 | 0.802 | <0.001 | 0.785 | <0.001 |
|  | Bimmen | -0.038 | 0.868 | <0.001 | 0.844 | <0.001 |
| Saar | Guedingen | 0.236 | 0.235 | 0.264 | 0.246 | 7.753 |
|  | Rehlingen | -0.128 | 0.5265 | 0.229 | 0.260 | 1.801 |
| Danube | Ulm | -0.132 | 0.580 | <0.001 | 0.960 | <0.001 |
|  | Jochenstein | -0.458 | 0.037 | 0.949 | 0.020 | 33.853 |

### Trends by trait categories

Except for the site Koblenz, all sites showed significant trends (Mann-Kendall) for at least one of the following indicators tested: Species richness, species proportion, or read proportion after grouping the fish into different groups (common, endangered, non-native, tolerant, sensitive, and migratory). Of the 78 tested cases (6 categories x 13 sites) across the sampling sites (arrows in figure 1), a total of 22 showed a significant decrease, 16 showed a significant increase and two showed conflicting significance. Common species (least concern) demonstrated a significantly declining trend at 46% of the sites (6) while increasing at only two sites (15%). Endangered species declined at 31% of sites (4) and increased at four sites (31%). Non-native species increased at 46% of sites (6) while decreasing only at one site (8%). Sensitive species declined at four sites (31%) and increased only at one site (8%). Tolerant species declined at three sites (23%). At two sites, conflicting significant trends were observed for tolerant species decreasing in species number but increasing at species proportion (Weil) or increasing in species proportion but decreasing in relative reads (Zehren). Migratory species decreased at four sites (31%) and increased at three (23%) (Figure 1 and Supplementary table 4).

### Correlation between relative reads and site occupancy trends

Our analysis revealed significant temporal changes for 25 fish species (40%), with 9 species showing significant trends in both occupancy and relative read abundance, 10 species showing changes only in occupancy, and 6 species only in relative abundance. Occupancy and relative read trends (Mann-Kendall tau) were generally correlated (rho = 0.56, p < 0.001) indicating that species expanding their range also increased in local read abundance.

Invasive *Neogobius melanostomus* demonstrated the strongest increases in both occupancy (τ = 0.759, p < 0.001) and relative reads (τ = 0.750, p < 0.001), followed by invasive *Babka gymnotrachelus* (occupancy; τ = 0.499, p = 0.020, relative reads; 0,472, p = 0.022) and native *Perca fluviatilis* (occupancy; τ = 0.706, p < 0.001, relative reads; 0.706, p < 0.001). Other species with significant occupancy or read abundance increases included *Chondrostoma nasus*, *Osmerus eperlanus*, *Phoxinus phoxinus*, and non-native *Ctenopharyngodon idella*. Conversely, several species declined significantly in one of the metrics including *Romanogobio* sp. (occupancy; τ = -0.574, p = 0.002), invasive *Ameiurus nebulosus* (occupancy τ = -0.596, p = 0.003; abundance τ = -0.486, p = 0.015), endangered *Lota lota* (occupancy; τ = -0.491, p = 0.011), *Vimba vimba* (occupancy; τ = -0.532, p = 0.006), *Abramis brama* (relative reads; τ = -0.529, p = 0.003) and *Barbatula barbatula* (occupancy; -0.471, p = 0.012) (Supplementary table 5).

The comparison of occupancy and read abundance trends revealed four distinct patterns in the correlation plot (Fig. 2): Quadrant I contains species with positive trends in both metrics, quadrant II includes species with declining occupancy but increasing relative abundance, quadrant III contains species with declining trends in both occupancy and abundance, and quadrant IV shows species with expanding occupancy but declining relative abundance, which contained only two species and generally weaker trends.

**Figure 2:**
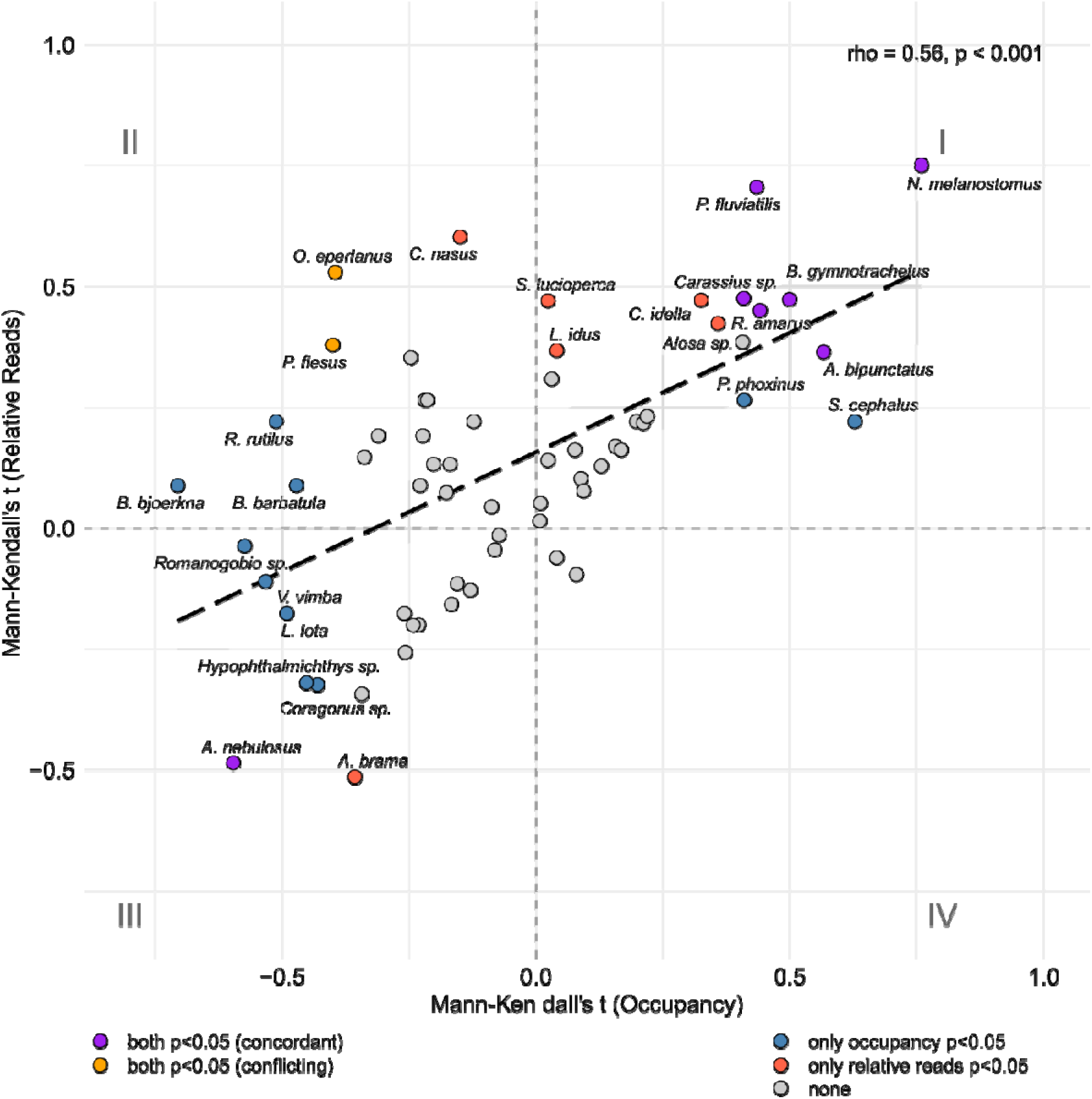
Mann-Kendall’s τ values for site occupancy (x-axis) and relative read abundance (y-axis) trends. Significant positive correlation (Spearman rho = 0.56, p < 0.001) indicates concordant in directional changes between the metrics. Colors indicate significance: purple = both metrics significant (p < 0.05), blue = occupancy only, orange = relative reads only, gray = neither significant.

### Spatiotemporal community change

Community composition within the same site changed steadily over time indicated both by significantly increasing Bray-Curtis dissimilarity and Jaccard dissimilarity (Figure 3 A, right). Analysis of temporal beta diversity within sites revealed that species replacement (turnover) was the primary driver of community change (0.314 ± 0.127). Nestedness accounted for the remaining portion (0.142 ± 0.035), indicating that while species replacement dominated, changes in species richness also contributed to temporal change. The analyses further revealed slightly increasing between-site dissimilarity, indicating that communities became less similar on average between sites. This was only significant using Bray-Curtis dissimilarity. Spatial variation between sites within each year was again dominated by species turnover (average: 0.482 ± 0.049) rather than nestedness (average: 0.163 ± 0.036) (Supplementary table 6).

**Figure 3:**
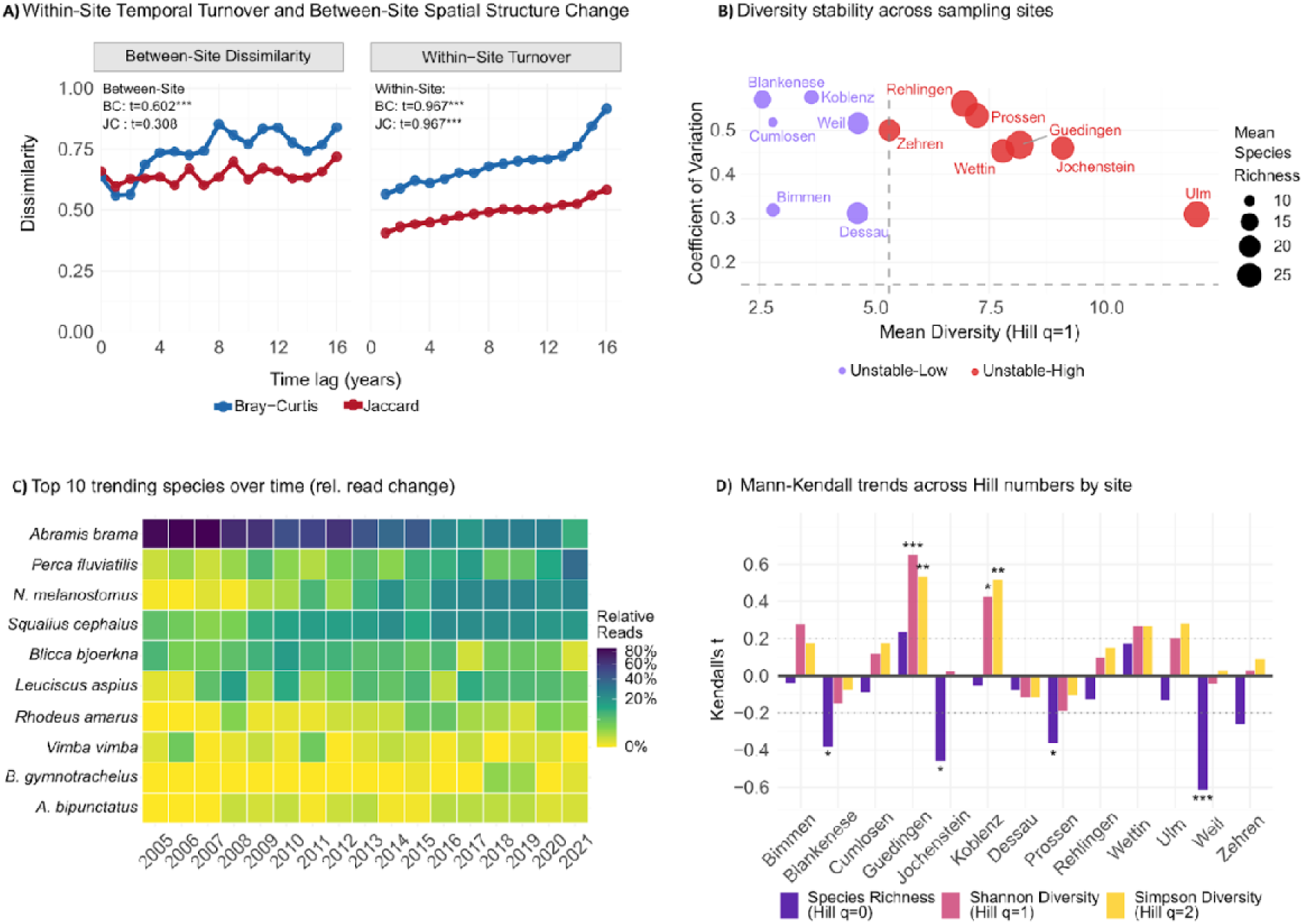
A) Beta diversity trends showing significant increases in both between-site spatial dissimilarity (left) and within-site temporal turnover (right) for Bray-Curtis (BC). Jaccard (JC) indices are only significant for within-site turnover. B) Sites by mean Shannon diversity and temporal variability (coefficient of variation), with point size representing species richness. Dashed lines indicate classification thresholds. Red = high diversity sites, blue = low diversity sites. C) Heatmap of relative read abundance for top 10 trending species across time. D) Alpha diversity trends per site inferred using Hill numbers with q=0: richness (p/a), q=1: Shannon, q=2: Simpson diversity. Kendall’s tau for correlation analysis results. Asterisks indicate significance (*p < 0.05, **p < 0.01, ***p < 0.001).

Per-site analyses revealed differences in both average diversity (q=1) and temporal stability (CV) across sites. Sites such as Ulm exhibited higher diversity and stability, whereas others, including Rehlingen, Blankenese and Koblenz, showed more variable and less diverse communities (Figure 3 B).

The heat map of the ten most trending species across sites (Figure 3 C) illustrates shifts in signal dominance through time. Early in the time-series *Abramis brama* was the most prominent taxon, but its relative abundance steadily declined. In contrast, the invasive round goby (*Neogobius melanostomus*) was increasingly strengthening in relative read abundance. Other species showed more site-specific or moderate changes.

Mann-Kendall trends across Hill numbers (q = 0, 1, 2) measuring richness, Shannon diversity, and Simpson diversity showed eleven sites decreasing in richness, with four sites significantly declining. Only Koblenz and Güdingen showed significant increases in Shannon and Simpson diversity over time (Figure 3 D), indicating increased community diversity, while other sites remained more stable.

### Fish species relative read abundance trends

Bayesian trend estimates of relative read abundance trends over time showed substantial species-specific variation. Most species (65%, n=41) showed stable read proportions over time. However, 12 species (19%) showed significantly negative trends, most pronounced in *Abramis brama* (estimate = -0.281), *Blicca bjoerkna* (estimate = -0.180), and *Barbatula barbatula* (estimate = -0.124). Ten species (16%) showed significantly positive trends, strongest in invasive *Neogobius melanostomus* (estimate = 0.457), followed by *Perca fluviatilis* (estimate = 0.210) and *Leuciscus aspius* (estimate = 0.120). Species classified as stable had moderate estimates with confidence intervals overlapping zero. The model diagnostics confirmed reliability with Rhat values at or near 1.0. Site occupancy over time was investigated using Spearman rank for three exemplary species. *Neogobius melanostomus* significantly increased in site occupancy (rho = 0.94), *Sander luciperca* remained stable (rho = 0.09) and *Blicca bjoerkna* significantly decreased (rho = -0.77) (Figure 4).

**Figure 4:**
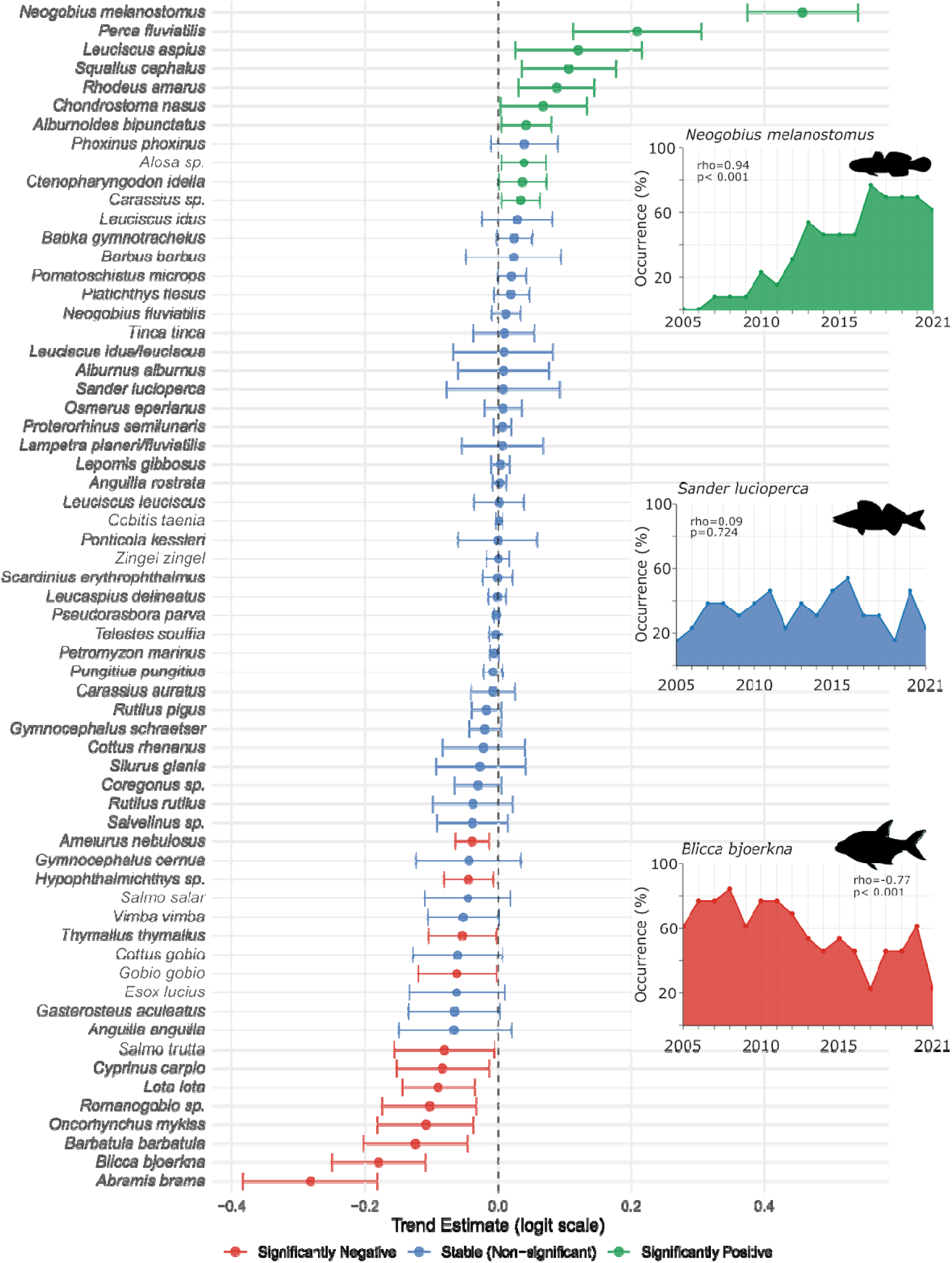
Temporal trends in fish species. The main panel (left) shows trend estimates (logit scale) based on Bayesian models with 95% confidence intervals for all species analyzed based on relative read abundance. Species are categorized as significantly positive (green), stable/non-significant (blue), or significantly negative (red). Right panels show exemplary three species: *Neogobius melanostomus* (top, green) showing a significant increase in site occurrence (rho=0.94, p<0.001), *Sander lucioperca* (middle, blue) exhibiting stable occurrence (rho=0.09, p=0.724), and *Blicca bjoerkna* (bottom, red) displaying a significant decline (rho=-0.77, p<0.001).

### Comparison of eDNA and traditional fish monitoring trend estimates

We compared the trends of 47 fish species that were identified in both datasets. The analysis revealed moderate to strong agreement across multiple statistical analysis despite the vast differences with respect to sampling design, frequency, method and data types.

Of the 47 species reported in both data sets, 35 (74.5%) showed directional trend agreement between the datasets. Spearman rank correlation was significant (rho = 0.550, p < 0.001), indicating moderate positive correlation between trend estimates (Figure 5 A). Cohen’s kappa coefficient was 0.471, indicating moderate agreement between the two datasets when trends were categorized. The Wilcoxon test showed no significant systematic bias of either dataset (p = 0.054) (Supplementary table 7). Bland-Altman analysis revealed a mean difference between dataset trend estimates of -0.043 with 95% limits of agreement ranging from -0.331 to 0.244. The relatively small mean difference and clustering of points around the zero line suggest good overall concordance between datasets, with only a few outlier species showing substantial disagreement (Figure 5 B). Aside from directional concordance, there are also several cases where the trend direction between datasets is slightly off due to estimates close to zero, however the delta between estimates is very small. An example for this is *Ponticola kessleri* with a delta of only 0.04 while it is increasing in the eDNA dataset (+0.0006) but decreasing in the regulatory monitoring data (-0.04). On the other hand there are species with concordant trend direction but a large delta, e.g., *Proterorhinus semilunaris* Δ = 0.64 (Figure 5, Supplement 7).

**Figure 5:**
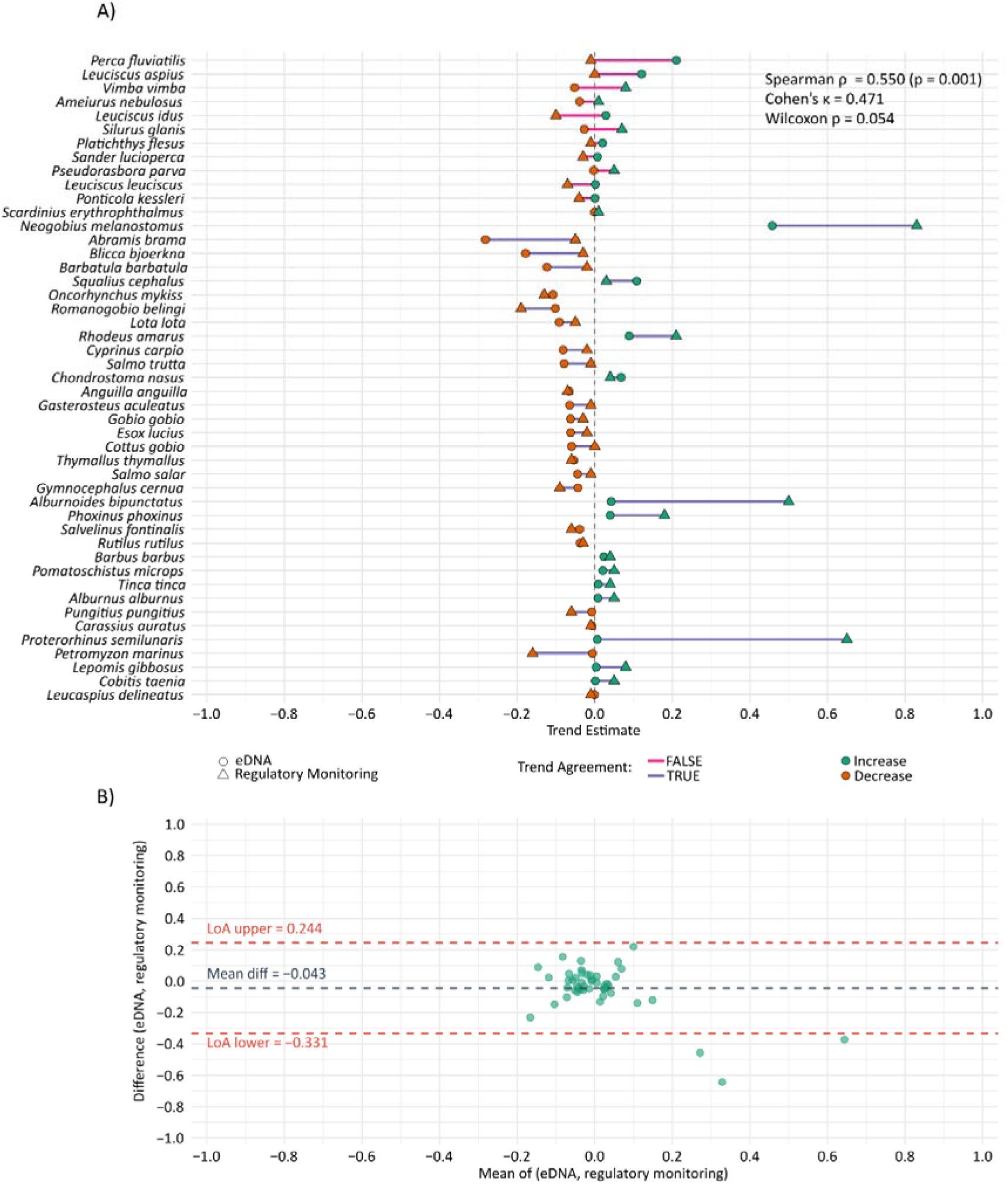
Comparison of fish species temporal trends between eDNA and regulatory monitoring datasets. A) Species-specific trend estimates from eDNA data (circles) and regulatory monitoring (triangles), color-coded by trend direction (orange = decrease, green = increase). Connecting lines indicate agreement between methods (blue = concordant trends, pink = discordant trends). B) Bland-Altman plot showing the agreement between dataset estimates. The mean difference between eDNA and regulatory monitoring trend estimates was -0.043 (black dashed line), with 95% limits of agreement ranging from -0.331 to 0.244 (red dashed lines).

### Physicochemical driver analysis

Of the 83 environmental parameters tested, 42 (51%) showed significant correlations with species richness (Supplementary table 8). Negative correlations dominated for contaminants and physical stressors (20 parameters, 24%), while positive correlations were observed primarily for nutrients and certain physical parameters (22 parameters, 26%). The strongest significant negative correlations were with discharge (rho = -0.62), organochlorine pesticides beta-HCH (rho = -0.58) and alpha-HCH (rho = -0.52), silt (rho = -0.56) (Figure 6 A), and chlorophyll a (rho = -0.47). Mercury, pH, water temperature, Lindane, or PFNA also negatively impacted richness estimates. Conversely, clay content (rho = 0.45), iron (rho = 0.40), and nutrients including phosphorus (rho = 0.38), nitrite (rho = 0.37), and nitrogen (rho = 0.36) showed positive associations with richness (Supplementary Table 8).

**Figure 6:**
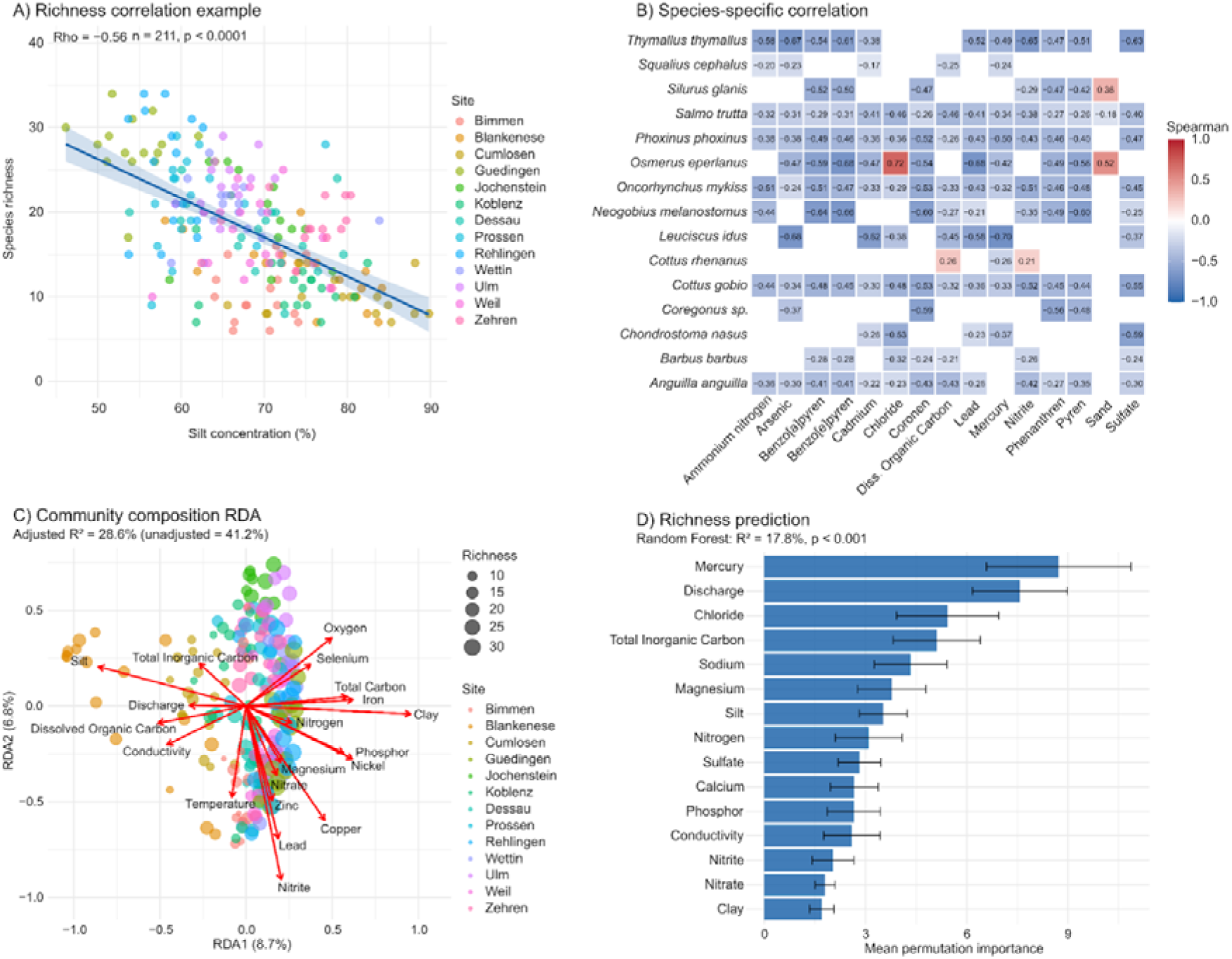
Analysis of fish community responses to environmental gradients. A) Species richness correlation with silt concentration as an example for a significant negative correlation (rho = -0.56, p < 0.0001). B) Heatmap of species-specific chemical sensitivities showing Spearman correlation coefficients between relative reads of 15 most impacted fish species and 15 most impactful environmental variables. Colors represent correlation strength and direction (red = positive, blue = negative), with only significant correlations (p < 0.05) displaying coefficient values. C) Redundancy analysis (RDA) showing community composition variation explained by chemical gradients (adjusted R² = 28.6%). Point size indicates species richness. Red arrows represent environmental variables with effect strength indicated by length, D) Random Forest prediction importance plot identifying key chemical predictors of species richness (R² = 17.8%, p < 0.001). Error bars indicate variability across model iterations.

The species-level analysis of 883 correlation tests revealed complex patterns based on relative read abundance (Figure 6 B, Supplementary table 8). 528 tests showed a negative correlation (59.8%) and 355 a positive (40.2%). Among the 82 parameters tested (one excluded), 81 (99%) showed negative associations with at least one species, 73 (89%) showed positive associations, and 72 (88%) exhibited both depending on the species tested. The parameters affecting the most species negatively were nitrite (14 species, mean rho = -0.37), ammonium nitrogen (13 species, mean rho = -0.36), and dissolved organic carbon (13 species, mean rho = - 0.35). Heavy metals also showed broad negative impacts with arsenic (11 species, mean rho = -0.40) and mercury (11 species, mean rho = -0.39). Polycyclic aromatic hydrocarbons had strong negative effects, with seven PAHs each affecting 8-11 species (mean rho = -0.44 to -0.50), led by Benzo(b)naphtho[2,1-d]thiophen (9 species, mean rho = -0.50). Legacy pesticides beta-HCH (10 species, mean rho = - 0.47) and alpha-HCH (7 species, mean rho = -0.50) also showed negative impacts. Oxygen concentration showed the broadest positive associations (11 species, mean rho = 0.32), followed by phosphorus (10 species, mean rho = 0.29) and iron (9 species, mean rho = 0.34).

Redundancy Analysis (RDA) explained 28.6% (41.2% unadjusted) of community compositional variation (adjusted R² = 0.286, p = 0.001, Figure 6 C). Sediment composition showed the strongest associations with clay (R² = 0.29) and silt (R² = 0.24). Nutrients (nitrite R² = 0.16, ammonium nitrogen R² = 0.09), heavy metals (nickel, iron, copper, lead; R² = 0.09-0.13), and oxygen (R² = 0.11) also contributed significantly. Partial redundancy analysis was used to separate the variation into spatial, temporal, and residual components. When controlling for spatial structure, the explained variance decreased to 5.5%, indicating 81% of the explained variance was associated with spatial differences. In contrast, conditioning on the year had little effect, suggesting temporally stable relationships. When both site and year were included as conditioning variables, 1.8% of community variation remained (adjusted R² = 0.018, p = 0.046), indicating a modest but statistically significant effect of the tested parameters independent of spatial or temporal composition (Supplementary table 8).

Random Forest analysis showed that environmental parameters significantly predicted species richness (R² = 0.178, RMSE = 6.2 species, MAE = 5.2 species, p < 0.001; Figure 6D). The 15 most influential predictors included mercury, discharge, chloride, and total inorganic carbon. Prediction accuracy varied among sites, ranging from Zehren (RMSE = 3.3 species) to Koblenz (RMSE = 9.4 species) (Supplementary table 8).

## Discussion

By analyzing archived SPM samples collected over 17 years from 13 sites across six major European rivers, we detected 63 fish species and reconstructed nearly two decades of community dynamics. Our data thus support earlier findings that eDNA-based biomonitoring from archived samples can reveal long-term ecological trends and serve as a powerful monitoring tool (Junk et al., 2025; Schütz et al., 2025; Sullivan et al., 2025).

Species detections were generally consistent with the expected fish fauna based on the German Red List, established monitoring programs, and regulatory monitoring, confirming both the reliability of our approach and the ecological plausibility of the results (Freyhof et al., 2023; ICPR, 2021; Igor Liška et al., 2021). River-specific detections, including Danube endemics (*Gymnocephalus schraetser*, *Rutilus pigus*) and coastal species (*Pomatoschistus microps*, *Osmerus eperlanus*) in estuaries, further demonstrate the method’s ability to reflect known biogeographic patterns. Although absolute species diversity was lower than the known total fish fauna in these rivers, this is expected given the limited spatial coverage of SPM sampling sites. The strong overlap with official records nevertheless demonstrates that eDNA from SPM provides an effective and reliable complement to conventional surveys, particularly for detecting standardized long-term trends.

### The eDNA SPM time series data (2005-2021) captures broader spatial and temporal trends in fish alpha and beta diversity from community trends to detailed patterns of key groups, as well as species-specific trends

Significant declines in fish species richness were observed at Jochenstein (Danube), Weil (Rhine), Blankenese (Elbe) and Prossen (Elbe), with no sites showing significant increases, raising concerns about environmental pressures reducing fish richness. However, decreasing species richness can indicate either ecosystem degradation or improvement in water quality through oligotrophication (Ibáñez et al., 2023). Many fish species thrive in the nutrient-rich conditions of the last decades, and restoration toward oligotrophic conditions may result in fewer but more specialized species reflecting natural reference communities (Kaijser et al., 2025). Recent analyses confirm that some species preferring unpolluted waters have shown positive trends in German rivers (Friedrichs-Manthey et al., 2024), while eutrophication can maintain high species richness of tolerant generalists (Kaijser et al., 2025). These patterns highlight that species richness alone is an insufficient metric. Investigating which species or functional groups increase and decrease is essential.

Grouping species by ecological traits and conservation status uncovered distinct, group-specific trends. Common species declined significantly at nearly half of the sites, indicating pressures on the baseline community or less favorable conditions for generalist species (e.g., through re-oligotrophication). Endangered species declined at four sites but increased at another four, suggesting strong site-specific variation in responses of vulnerable taxa.

Non-native species, in contrast, increased significantly at almost half of the sites, reflecting the continued expansion of non-native fishes observed in European rivers (Leuven et al., 2009; Zieritz et al., 2017). Sensitive species declined at four sites and increased at only one, supporting concerns about the loss of pollution- and habitat-sensitive taxa (Schinegger et al., 2012). Migratory species exhibited mixed patterns, likely reflecting site-specific differences in connectivity or restoration status (Jähnig et al., 2011). Conflicting results for tolerant species declining in richness but increasing in proportional abundance at some sites may indicate shifts in dominance structure, stocking, or reflect limitations of eDNA relative read-based inference.

A key limitation of eDNA metabarcoding remains its semi-quantitative nature, as read counts do not directly correspond to biomass or abundance (Rourke et al., 2022). To evaluate semi-quantitative patterns, we compared relative read abundance and site occupancy, which showed significantly correlated temporal trends across species, supporting previous work linking eDNA signal strength to fish abundance (Lacoursière-Roussel et al., 2016; Yates et al., 2019). Species generally exhibited concurrent increases or decreases in both metrics, such as rising trends in invasive gobiids (*N. melanostomus*, *Babka gymnotrachelus*) and declines in native cyprinids and *L. lota*. Evaluating both metrics enhances the ecological interpretability of eDNA data, complementing qualitative detection with semi-quantitative information.

Community compositions changed substantially over the 17-year period, reflected in both temporal turnover within sites and spatial differentiation among sites. Within-site temporal turnover increased significantly with time for both Bray-Curtis and Jaccard dissimilarities, indicating compositional change driven primarily by species replacement rather than nestedness. This pattern is consistent with species reassembly under environmental change (Dornelas et al., 2013) rather than simple species loss. Simultaneously, between-site dissimilarity increased over the study period for Bray-Curtis suggesting community differentiation rather than biotic homogenization (Olden & Rooney, 2006; Rolls et al., 2023). Shifts in site-specific dominance such as the decline in relative read abundance of *Abramis brama* and expansion of invasive *Neogobius melanostomus*, illustrate these altered communities (Brandner et al., 2013). These patterns generally align with global trends of freshwater community reorganization under environmental change (Blowes et al., 2019; Magurran et al., 2019). Local extinctions and introductions caused by e.g., invasive species or local restocking and reintroduction programs drive temporal turnover and promote spatial differentiation (Magalhães et al., 2020; Rolls et al., 2023).

Temporal responses differed between sites. All 13 sites were classified as either “Unstable-High” (n=7) or “Unstable-Low” (n=6), with no sites achieving “Stable-High” conditions. Mann-Kendall trend analyses across Hill numbers revealed declining richness at 11 sites, with four showing significant losses. Only Koblenz and Güdingen showed significant increases in both Shannon and Simpson diversity. These results indicate that local environmental stability supports community persistence (Gonzalez & Loreau, 2009), and that sites exhibiting high variability or declining diversity are in need of targeted investigation and management interventions (Birk et al., 2020).

Interpreting eDNA signals for commercially stocked taxa (e.g., *Oncorhynchus mykiss*, *Salmo salar*., etc.) also remains challenging, as stocking and fisheries management can mask natural patterns. Despite these uncertainties, most of the 63 detected taxa remained stable, while a substantial subset showed significant directional changes aligning with recent assessments of German fish communities (Friedrichs-Manthey et al., 2024). Although we do not compare our findings with data from before 2005, this also shows that freshwater recovery has largely stagnated in recent years (Haase et al., 2023) or continues to decline at certain sites and for certain groups and species. Declines in sensitive species such as *Barbatula barbatula* are concerning, as these taxa are often affected by habitat degradation, altered flow, and sedimentation (Radinger & Wolter, 2015; Schinegger et al., 2012). Conversely, declines in widespread eutrophic species such as *Abramis brama* or *Blicca bjoerkna* may signal re-oligotrophication through restoration efforts and water quality improvement. Decreases in cold-water species such as *Salmo trutta* and *Lota lota* further highlight their sensitivity to warming and hydromorphological alteration (Comte et al., 2013; Filipe et al., 2013).

Several species showed strong positive trends. The invasive round goby (*Neogobius melanostomus*) increased steeply, consistent with its rapid expansion across Central Europe and concurrent declines of native benthic fishes such as *Barbatua barbatula* and *Gobio gobio* (Błońska et al., 2024; Lampert et al., 2024), for which a decline was also observed in our study. Increases in tolerant species, including *Perca fluviatilis* and *Leuciscus aspius*, likely reflect their adaptability to altered habitats and nutrient enrichment (Friedrichs-Manthey et al., 2024). Positive trends in additional non-native and fishery-relevant taxa (e.g., *Ctenopharyngodon idella*) further demonstrate how introductions continue to reshape community composition (Haubrock et al., 2022).

While many taxa remained stable, declines in sensitive and specialist species alongside increases in tolerant and invasive taxa indicate an ongoing shift in community structure. The observed declines in some endangered or sensitive species are particularly concerning, emphasizing the need for proactive conservation and habitat restoration under the EU Water Framework Directive or the Nature Restoration Regulation, as well as improved control of invasive species.

### Spatiotemporal patterns of fish biodiversity change identified with the eDNA metabarcoding dataset generally align with those observed in regulatory fish monitoring

Comparison with regulatory monitoring showed a broad agreement between the two datasets. Both approaches captured similar population trends, with nearly three-quarters of the shared species showing a directional agreement. Previous studies likewise showed that eDNA and conventional surveys generate largely similar results (Czeglédi et al., 2021; Wang et al., 2024). However, directional agreement alone does not fully capture concordance as the magnitude of differences varied substantially among species. For instance, *Proterorhinus semilunaris* and *Alburnoides bipunctatus* showed increasing trends in both datasets yet differed by 0.64 and 0.46 trend units, far exceeding typical effect sizes. Conversely, some directionally discordant species showed remarkably close estimates such as *Ponticola kessleri* (Δ = 0.04) and Ameiurus nebulosus (Δ = 0.05) which differed primarily in whether marginal trends near zero were classified as slight increases versus slight decreases. Cohen’s kappa and the Wilcoxon test suggest the absence of systematic bias or the underestimation of trends relative to the regulatory monitoring dataset. The Bland-Altman analysis further supports this, with relatively narrow limits of agreement and clustering around zero, indicating that discrepancies are limited to a few outlier species. Such differences likely reflect species-specific detection biases, differences in life-history, or habitat use that affect capture probability in nets versus DNA shedding rates in SPM samples (Curto et al., 2025; Rourke et al., 2022; Thalinger et al., 2021).

Despite substantial differences in sampling design, effort, and frequency, eDNA and regulatory monitoring datasets showed concordant trends for the majority of species. eDNA from SPM provides a broad taxonomic coverage with high temporal resolution from existing archives, whereas conventional surveys provide actual abundance data from more sites but with lower temporal resolution. Integrating both approaches can therefore strengthen long-term biodiversity assessments, with eDNA time series complementing conventional monitoring to improve confidence in trend detection across freshwater fish communities at a higher temporal resolution and a lower cost.

### Physiochemical environmental parameters obtained majorly from the same SPM samples explain a significant variation in fish species richness and community composition, with distinct effects of pollutants, nutrients, and physical gradients

Multiple environmental stressors shaped fish richness and diversity at the sampling sites. Negative correlations with contaminants like pesticides and heavy metals, physical stressors like temperature and conductivity, and fine sediments underline the effects of habitat degradation and pollution on freshwater assemblages (Feio et al., 2023; Walsh et al., 2005). Concerningly, higher temperatures were also significantly correlated to lower species richness, raising concerns for the future under the current climate change. Conversely, higher oxygen, moderate nutrients and iron appeared to support higher richness, possibly reflecting higher productivity when nutrient levels remain below toxic eutrophication thresholds (Horka et al., 2023).

The strong negative correlation between richness and silt content highlights the ecological importance of fine sediment (do Amaral et al., 2024; Kemp et al., 2011). Blankenese in the Elbe delta, which experienced declining species richness, also showed the highest and increasing fine sediment concentrations. A recent study linked declining fish diversity in the Elbe delta partially to the high fine sediment content caused by dredging for shipping in recent years (Theilen et al., 2025). Although this negative correlation between fish richness and fine sediment content is well supported, further tests should be carried out to investigate if the sediment composition has an effect on the eDNA detection probability due to binding efficiency (Brandão-Dias et al., 2023; Díaz et al., 2020) or inhibition (Buxton et al., 2017; Goldberg et al., 2016).

Species-specific responses emphasized that not a single factor drives community change but rather, a mosaic of multiple stressors leads to cumulative or sometimes opposing influences (Kaijser et al., 2025; Madge Pimentel et al., 2025; Vos et al., 2023). Widespread negative associations with heavy metals, legacy pesticides, and polycyclic aromatic hydrocarbons are consistent with their persistent toxicity in aquatic ecosystems, while positive links with e.g., oxygen likely indicate healthier conditions (Malaj et al., 2014; van der Oost et al., 2003). Invasive *Neogobius melanostomus* showed a positive correlation with warmer temperatures, raising concerns about its future spread under climate change.

Redundancy analysis indicated that environmental gradients, particularly sediment and metal concentrations, explained about 29% community composition variation. However, most of this signal reflected spatial rather than temporal differences, suggesting site-specific but temporally stable stressors. After accounting for space and time, a small but significant residual effect persisted, indicating that the measured environmental parameters continue to shape local fish communities independent of spatial and temporal structure.

Random forest modeling reinforced that fish richness responds to interacting stressors rather than single dominant factors with mercury, discharge, chloride, and several nutrients as key predictors. This underscores the need for integrated management addressing both chemical pollution and hydrological alteration (Birk et al., 2020; Ormerod et al., 2010). However, the unexplained portion suggests that additional factors including more environmental parameters, habitat structure, land use, habitat degradation, and biotic interactions need further investigation. Particularly, higher temporal resolution of both biodiversity data (from eDNA metabarcoding or other methods) and underlying physico-chemical data are essential to increase the explanatory power of driver analysis in a multiple stressor framework.

## Conclusion

Our comprehensive 17-year analysis demonstrates that eDNA metabarcoding of archived SPM samples provides a powerful and reliable tool for long-term biodiversity assessment in river ecosystems. The approach enables simultaneous reconstruction of biological trends and physiochemical stressors from the same standardized samples, integrating biodiversity and pollution assessment within a single monitoring framework. This supports a One Health and early-warning perspective for ecosystem monitoring and recovery assessment.

The study shows that fish community dynamics, including species invasions, and range shifts, can be reconstructed retrospectively and tracked through time, providing temporal resolution comparable to, and largely congruent with, regulatory monitoring programs covering thousands of sites. Archived SPM thus can serve as a near “all-in-one” monitoring matrix, bridging molecular biodiversity assessment and chemical analyses within environmental specimen banks.

Expanding this SPM approach to additional taxonomic groups will enable comprehensive tree-of-life ecosystem monitoring linked to physiochemical parameters. This integrated framework bridges chemical and biological monitoring, supporting evidence-based management under global biodiversity conservation and environmental protection initiatives.

## Supporting information

Taxa_final

Grouptrends

Read_Occupancy_Correlation

Betadiversity

Trend_Agreement

Chem_Correlation

parameters

taxa-raw

## References

1. Altermatt, F., Couton, M., Carraro, L., Keck, F., Lawson-Handley, L., Leese, F., Zhang, X., Zhang, Y., & Blackman, R. C. (2025). Utilizing aquatic environmental DNA to address global biodiversity targets. Nature Reviews Biodiversity, 1–15. 10.1038/s44358-025-00044-x

2. Beng, K. C., & Corlett, R. T. (2020). Applications of environmental DNA (eDNA) in ecology and conservation: Opportunities, challenges and prospects. Biodiversity and Conservation, 29(7), 2089–2121. 10.1007/s10531-020-01980-0

3. Birk, S., Bonne, W., Borja, A., Brucet, S., Courrat, A., Poikane, S., Solimini, A., van de Bund, W., Zampoukas, N., & Hering, D. (2012). Three hundred ways to assess Europe’s surface waters: An almost complete overview of biological methods to implement the Water Framework Directive. Ecological Indicators, 18, 31–41. 10.1016/j.ecolind.2011.10.009

4. Birk, S., Chapman, D., Carvalho, L., Spears, B. M., Andersen, H. E., Argillier, C., Auer, S., Baattrup-Pedersen, A., Banin, L., Beklioğlu, M., Bondar-Kunze, E., Borja, A., Branco, P., Bucak, T., Buijse, A. D., Cardoso, A. C., Couture, R.-M., Cremona, F., de Zwart, D., … Hering, D. (2020). Impacts of multiple stressors on freshwater biota across spatial scales and ecosystems. Nature Ecology & Evolution, 4(8), 1060–1068. 10.1038/s41559-020-1216-4

5. Bista, I., Carvalho, G. R., Walsh, K., Seymour, M., Hajibabaei, M., Lallias, D., Christmas, M., & Creer, S. (2017). Annual time-series analysis of aqueous eDNA reveals ecologically relevant dynamics of lake ecosystem biodiversity. Nature Communications, 8(1), 14087. 10.1038/ncomms14087

6. Błońska, D., Janic, B., Tarkan, A. S., Piria, M., Bănăduc, D., Švolíková, K. S., Števove, B., Lappalainen, J., Pyrzanowski, K., Tszydel, M., & Bukowska, B. (2024). Physiological responses of invasive round goby (Neogobius melanostomus) to environmental stressors across a latitudinal span. Biological Invasions, 26(10), 3433–3444. 10.1007/s10530-024-03387-2

7. Blowes, S. A., Supp, S. R., Antão, L. H., Bates, A., Bruelheide, H., Chase, J. M., Moyes, F., Magurran, A., McGill, B., Myers-Smith, I. H., Winter, M., Bjorkman, A. D., Bowler, D. E., Byrnes, J. E. K., Gonzalez, A., Hines, J., Isbell, F., Jones, H. P., Navarro, L. M., … Dornelas, M. (2019). The geography of biodiversity change in marine and terrestrial assemblages. Science. 10.1126/science.aaw1620

8. Bohmann, K., Elbrecht, V., Carøe, C., Bista, I., Leese, F., Bunce, M., Yu, D. W., Seymour, M., Dumbrell, A. J., & Creer, S. (2022). Strategies for sample labelling and library preparation in DNA metabarcoding studies. Molecular Ecology Resources, 22(4), 1231–1246. 10.1111/1755-0998.13512

9. Brandão-Dias, P. F. P., Tank, J. L., Snyder, E. D., Mahl, U. H., Peters, B., Bolster, D., Shogren, A. J., Lamberti, G. A., Bibby, K., & Egan, S. P. (2023). Suspended Materials Affect Particle Size Distribution and Removal of Environmental DNA in Flowing Waters. Environmental Science & Technology, 57(35), 13161–13171. 10.1021/acs.est.3c02638

10. Brandner, J., Cerwenka, A. F., Schliewen, U. K., & Geist, J. (2013). Bigger Is Better: Characteristics of Round Gobies Forming an Invasion Front in the Danube River. PLOS ONE, 8(9), e73036. 10.1371/journal.pone.0073036

11. Buchner, D. (2022). Guanidine-based DNA extraction with silica-coated beads or silica spin columns. https://www.protocols.io/view/guanidine-based-dna-extraction-with-silica-coated-che7t3hn

12. Buchner, D., Macher, T.-H., & Leese, F. (2022). APSCALE: Advanced pipeline for simple yet comprehensive analyses of DNA metabarcoding data. Bioinformatics, 38(20), 4817–4819. 10.1093/bioinformatics/btac588

13. Buxton, A. S., Groombridge, J. J., Zakaria, N. B., & Griffiths, R. A. (2017). Seasonal variation in environmental DNA in relation to population size and environmental factors. Scientific Reports, 7(1), 46294. 10.1038/srep46294

14. Carraro, L., Mächler, E., Wüthrich, R., & Altermatt, F. (2020). Environmental DNA allows upscaling spatial patterns of biodiversity in freshwater ecosystems. Nature Communications, 11(1), 3585. 10.1038/s41467-020-17337-8

15. Comte, L., Buisson, L., Daufresne, M., & Grenouillet, G. (2013). Climate-induced changes in the distribution of freshwater fish: Observed and predicted trends. Freshwater Biology, 58(4), 625–639. 10.1111/fwb.12081

16. Curto, M., Batista, S., Santos, C. D., Ribeiro, F., Nogueira, S., Ribeiro, D., Prindle, B., Licari, D., Riccioni, G., Dias, D., Pina-Martins, F., Jentoft, S., Veríssimo, A., Alves, M. J., & Gante, H. F. (2025). Freshwater fish community assessment using eDNA metabarcoding vs. capture-based methods: Differences in efficiency and resolution coupled to habitat and ecology. Environmental Research, 274, 121238. 10.1016/j.envres.2025.121238

17. Czeglédi, I., Sály, P., Specziár, A., Preiszner, B., Szalóky, Z., Maroda, Á., Pont, D., Meulenbroek, P., Valentini, A., & Erős, T. (2021). Congruency between two traditional and eDNA-based sampling methods in characterising taxonomic and trait-based structure of fish communities and community-environment relationships in lentic environment. Ecological Indicators, 129, 107952. 10.1016/j.ecolind.2021.107952

18. Díaz, C., Wege, F.-F., Tang, C. Q., Crampton-Platt, A., Rüdel, H., Eilebrecht, E., & Koschorreck, J. (2020). Aquatic suspended particulate matter as source of eDNA for fish metabarcoding. Scientific Reports, 10(1), 14352. 10.1038/s41598-020-71238-w

19. Djurhuus, A., Closek, C. J., Kelly, R. P., Pitz, K. J., Michisaki, R. P., Starks, H. A., Walz, K. R., Andruszkiewicz, E. A., Olesin, E., Hubbard, K., Montes, E., Otis, D., Muller-Karger, F. E., Chavez, F. P., Boehm, A. B., & Breitbart, M. (2020). Environmental DNA reveals seasonal shifts and potential interactions in a marine community. Nature Communications, 11(1), 254. 10.1038/s41467-019-14105-1

20. do Amaral, P. H. M., Linares, M. S., de Oliveira Tourinho, T. C., Hughes, R. M., & Callisto, M. (2024). Fine sediments produce tipping points in the taxonomic and functional structure of benthic macroinvertebrates in neotropical streams. Aquatic Sciences, 87(1), 17. 10.1007/s00027-024-01144-0

21. Dornelas, M., Magurran, A. E., Buckland, S. T., Chao, A., Chazdon, R. L., Colwell, R. K., Curtis, T., Gaston, K. J., Gotelli, N. J., Kosnik, M. A., McGill, B., McCune, J. L., Morlon, H., Mumby, P. J., Øvreås, L., Studeny, A., & Vellend, M. (2013). Quantifying temporal change in biodiversity: Challenges and opportunities. Proceedings of the Royal Society B: Biological Sciences, 280(1750), 20121931. 10.1098/rspb.2012.1931

22. Feio, M. J., Hughes, R. M., Serra, S. R. Q., Nichols, S. J., Kefford, B. J., Lintermans, M., Robinson, W., Odume, O. N., Callisto, M., Macedo, D. R., Harding, J. S., Yates, A. G., Monk, W., Nakamura, K., Mori, T., Sueyoshi, M., Mercado-Silva, N., Chen, K., Baek, M. J., … Sharma, S. (2023). Fish and macroinvertebrate assemblages reveal extensive degradation of the world’s rivers. Global Change Biology, 29(2), 355–374. 10.1111/gcb.16439

23. Filipe, A. F., Lawrence, J. E., & Bonada, N. (2013). Vulnerability of stream biota to climate change in mediterranean climate regions: A synthesis of ecological responses and conservation challenges. Hydrobiologia, 719(1), 331–351. 10.1007/s10750-012-1244-4

24. Freyhof, J.; Bowler, D.; Broghammer, T.; Friedrichs-Manthey, M.; Heinze, S. & Wolter, C. (2023). Rote Liste und Gesamtartenliste der sich im Süßwasser reproduzierenden Fische und Neunaugen (Pisces et Cyclostomata) Deutschlands. Landwirtschaftsverlag GmbH. 10.19213/972176

25. Friedrichs-Manthey, M., Bowler, D. E., & Freyhof, J. (2024). Freshwater fish in mid and northern German rivers – Long-term trends and associated species traits. Science of The Total Environment, 957, 177759. 10.1016/j.scitotenv.2024.177759

26. Gaye-Siessegger, J., Bader, S., Haberbosch, R., & Brinker, A. (2022). Spread of invasive Ponto-Caspian gobies and their effect on native fish species in the Neckar River (South Germany). https://kops.uni-konstanz.de/handle/123456789/59001

27. Goldberg, C. S., Turner, C. R., Deiner, K., Klymus, K. E., Thomsen, P. F., Murphy, M. A., Spear, S. F., McKee, A., Oyler-McCance, S. J., Cornman, R. S., Laramie, M. B., Mahon, A. R., Lance, R. F., Pilliod, D. S., Strickler, K. M., Waits, L. P., Fremier, A. K., Takahara, T., Herder, J. E., & Taberlet, P. (2016). Critical considerations for the application of environmental DNA methods to detect aquatic species. Methods in Ecology and Evolution, 7(11), 1299–1307. 10.1111/2041-210X.12595

28. Gonzalez, A., & Loreau, M. (2009). The Causes and Consequences of Compensatory Dynamics in Ecological Communities. Annual Review of Ecology, Evolution, and Systematics, 40(Volume 40, 2009), 393–414. 10.1146/annurev.ecolsys.39.110707.173349

29. Grabowska, J., Błońska, D., Ondračková, M., & Kakareko, T. (2023). The functional ecology of four invasive Ponto–Caspian gobies. Reviews in Fish Biology and Fisheries, 33(4), 1329–1352. 10.1007/s11160-023-09801-7

30. Haase, P., Bowler, D. E., Baker, N. J., Bonada, N., Domisch, S., Garcia Marquez, J. R., Heino, J., Hering, D., Jähnig, S. C., Schmidt-Kloiber, A., Stubbington, R., Altermatt, F., Álvarez-Cabria, M., Amatulli, G., Angeler, D. G., Archambaud-Suard, G., Jorrín, I. A., Aspin, T., Azpiroz, I., … Welti, E. A. R. (2023). The recovery of European freshwater biodiversity has come to a halt. Nature, 620(7974), 582–588. 10.1038/s41586-023-06400-1

31. Haase, P., Tonkin, J. D., Stoll, S., Burkhard, B., Frenzel, M., Geijzendorffer, I. R., Häuser, C., Klotz, S., Kühn, I., McDowell, W. H., Mirtl, M., Müller, F., Musche, M., Penner, J., Zacharias, S., & Schmeller, D. S. (2018). The next generation of site-based long-term ecological monitoring: Linking essential biodiversity variables and ecosystem integrity. Science of The Total Environment, 613–614, 1376–1384. 10.1016/j.scitotenv.2017.08.111

32. Haubrock, P. J., Ahmed, D. A., Cuthbert, R. N., Stubbington, R., Domisch, S., Marquez, J. R. G., Beidas, A., Amatulli, G., Kiesel, J., Shen, L. Q., Soto, I., Angeler, D. G., Bonada, N., Cañedo-Argüelles, M., Csabai, Z., Datry, T., de Eyto, E., Dohet, A., Drohan, E., … Haase, P. (2022). Invasion impacts and dynamics of a European-wide introduced species. Global Change Biology, 28(15), 4620–4632. 10.1111/gcb.16207

33. Horka, P., Musilova, Z., Holubova, K., Jandova, K., Kukla, J., Rutkayova, J., & Jones, J. I. (2023). Anthropogenic nutrient loading affects both individual species and the trophic structure of river fish communities. Frontiers in Ecology and Evolution, 10. 10.3389/fevo.2022.1076451

34. Hupalo, K., Macher, T.-H., Schütz, R., & Leese, F. (2022). Assessing Metropolitan Biodiversity Using Aquatic Environmental DNA Metabarcoding. In J. M. Gurr, R. Parr, & D. Hardt (Eds.), Urban Studies (1st ed., pp. 223–248). transcript Verlag. 10.14361/9783839463109-013

35. Ibáñez, C., Caiola, N., Barquín, J., Belmar, O., Benito-Granell, X., Casals, F., Fennessy, S., Hughes, J., Palmer, M., Peñuelas, J., Romero, E., Sardans, J., & Williams, M. (2023). Ecosystem-level effects of re-oligotrophication and N:P imbalances in rivers and estuaries on a global scale. Global Change Biology, 29(5), 1248–1266. 10.1111/gcb.16520

36. ICPR, I. (2021). Fish in the Rhine 2018/2019. https://www.iksr.org/fileadmin/user_upload/DKDM/Dokumente/Fachberichte/DE/rp_De_0279.pdf

37. Igor Liška, Franz Wagner, Manfred Sengl, Karin Deutsch, & Jaroslav Slobodník and Momir Paunović. (2021). JDS 4. https://www.danubesurvey.org/jds4/publications/scientific-report

38. Jähnig, S. C., Lorenz, A. W., Hering, D., Antons, C., Sundermann, A., Jedicke, E., & Haase, P. (2011). River restoration success: A question of perception. Ecological Applications, 21(6), 2007–2015.

39. Junk, I., Hans, J., Perez-Lamarque, B., Stothut, M., Weber, S., Gold, E., Schubert, C., Schumacher, A., Schmitt, N., Melcher, A., Paulus, M., Klein, R., Teubner, D., Koschorreck, J., Kennedy, S., Morlon, H., & Krehenwinkel, H. (2025). Archived natural DNA samplers reveal four decades of biodiversity change across the tree of life. Nature Ecology & Evolution, 9(10), 1873–1884. 10.1038/s41559-025-02812-6

40. Kaijser, W., Musiol, M., Schneider, A. R., Prati, S., Brauer, V. S., Bayer, R., Birk, S., Brauns, M., Dunne, L., Enss, J., Farias, L., Feld, C. K., Feldhaus, L., Gillmann, S. M., Hupało, K., Osakpolor, S. E., Olberg, S. L. M., Pimentel, I. M., Schäfer, R. B., … Hering, D. (2025). Meta-analysis-derived estimates of stressor–response associations for riverine organism groups. Nature Ecology & Evolution, 9(12), 2304–2321. 10.1038/s41559-025-02884-4

41. Kemp, P., Sear, D., Collins, A., Naden, P., & Jones, I. (2011). The impacts of fine sediment on riverine fish. Hydrological Processes, 25(11), 1800–1821. 10.1002/hyp.7940

42. Krehenwinkel, H., Weber, S., Broekmann, R., Melcher, A., Hans, J., Wolf, R., Hochkirch, A., Kennedy, S. R., Koschorreck, J., Künzel, S., Müller, C., Retzlaff, R., Teubner, D., Schanzer, S., Klein, R., Paulus, M., Udelhoven, T., & Veith, M. (2022). Environmental DNA from archived leaves reveals widespread temporal turnover and biotic homogenization in forest arthropod communities. eLife, 11, e78521. 10.7554/eLife.78521

43. Lacoursière-Roussel, A., Rosabal, M., & Bernatchez, L. (2016). Estimating fish abundance and biomass from eDNA concentrations: Variability among capture methods and environmental conditions. Molecular Ecology Resources, 16(6), 1401–1414. 10.1111/1755-0998.12522

44. Lampert, K. P., Heermann, L., Storm, S., Hirsch, P. E., Cerwenka, A. F., Heubel, K., Borcherding, J., & Waldvogel, A.-M. (2024). Round gobies (Neogobius melanostomus) in the River Rhine: Population genetic support for invasion via two different routes. PLOS ONE, 19(9), e0310692. 10.1371/journal.pone.0310692

45. Leray, M., Knowlton, N., & Machida, R. J. (2022). MIDORI2: A collection of quality controlled, preformatted, and regularly updated reference databases for taxonomic assignment of eukaryotic mitochondrial sequences. Environmental DNA, 4(4), 894–907. 10.1002/edn3.303

46. Leuven, R. S. E. W., van der Velde, G., Baijens, I., Snijders, J., van der Zwart, C., Lenders, H. J. R., & bij de Vaate, A. (2009). The river Rhine: A global highway for dispersal of aquatic invasive species. Biological Invasions, 11(9), 1989–2008. 10.1007/s10530-009-9491-7

47. Liang, D., Xia, J., Song, J., Sun, H., & Xu, W. (2022). Using eDNA to Identify the Dynamic Evolution of Multi-Trophic Communities Under the Eco-Hydrological Changes in River. Frontiers in Environmental Science, 10. 10.3389/fenvs.2022.929541

48. Lothian, A. J., Bolland, J. D., Albright, A. J., Jubb, W. M., Bubb, D. H., Noble, R. A. A., Nunn, A. D., Dodd, J. R., Tummers, J. S., & Lucas, M. C. (2024). Factors influencing European river lamprey passage at a tidal river barrier. Hydrobiologia, 851(20), 4803–4820. 10.1007/s10750-024-05633-z

49. Macher, T.-H., Beermann, A. J., & Leese, F. (2021). TaxonTableTools: A comprehensive, platform-independent graphical user interface software to explore and visualise DNA metabarcoding data. Molecular Ecology Resources, 21(5), 1705–1714. 10.1111/1755-0998.13358

50. Macher, T.-H., Schütz, R., Arle, J., Beermann, A. J., Koschorreck, J., & Leese, F. (2021). Beyond fish eDNA metabarcoding: Field replicates disproportionately improve the detection of stream associated vertebrate species. Metabarcoding and Metagenomics, 5, e66557. 10.3897/mbmg.5.66557

51. Macher, T.-H., Schütz, R., Yildiz, A., Beermann, A. J., & Leese, F. (2023). Evaluating five primer pairs for environmental DNA metabarcoding of Central European fish species based on mock communities. Metabarcoding and Metagenomics, 7, e103856. 10.3897/mbmg.7.103856

52. Madge Pimentel, I., Albini, D., Beermann, A. J., Leese, F., Macaulay, S. J., Matthaei, C. D., Orr, J. A., Piggott, J. J., & Schäfer, R. B. (2025). Hypothesis-Driven Research on Multiple Stressors: An Analytical Framework for Stressor Interactions. Ecology and Evolution, 15(8), e71959. 10.1002/ece3.71959

53. Magalhães, A. L. B., Daga, V. S., Bezerra, L. A. V., Vitule, J. R. S., Jacobi, C. M., & Silva, L. G. M. (2020). All the colors of the world: Biotic homogenization-differentiation dynamics of freshwater fish communities on demand of the Brazilian aquarium trade. Hydrobiologia, 847(18), 3897–3915. 10.1007/s10750-020-04307-w

54. Magurran, A. E., Dornelas, M., Moyes, F., & Henderson, P. A. (2019). Temporal β diversity—A macroecological perspective. Global Ecology and Biogeography, 28(12), 1949–1960. 10.1111/geb.13026

55. Malaj, E., von der Ohe, P. C., Grote, M., Kühne, R., Mondy, C. P., Usseglio-Polatera, P., Brack, W., & Schäfer, R. B. (2014). Organic chemicals jeopardize the health of freshwater ecosystems on the continental scale. Proceedings of the National Academy of Sciences, 111(26), 9549–9554. 10.1073/pnas.1321082111

56. Olden, J. D., & Rooney, T. P. (2006). On defining and quantifying biotic homogenization. Global Ecology and Biogeography, 15(2), 113–120. 10.1111/j.1466-822X.2006.00214.x

57. Ormerod, S. J., Dobson, M., Hildrew, A. G., & Townsend, C. R. (2010). Multiple stressors in freshwater ecosystems. Freshwater Biology, 55(s1), 1–4. 10.1111/j.1365-2427.2009.02395.x

58. Radinger, J., Britton, J. R., Carlson, S. M., Magurran, A. E., Alcaraz-Hernández, J. D., Almodóvar, A., Benejam, L., Fernández-Delgado, C., Nicola, G. G., Oliva-Paterna, F.J., Torralva, M., & García-Berthou, E. (2019). Effective monitoring of freshwater fish. Fish and Fisheries, 20(4), 729–747. 10.1111/faf.12373

59. Radinger, J., & Wolter, C. (2015). Disentangling the effects of habitat suitability, dispersal, and fragmentation on the distribution of river fishes. Ecological Applications, 25(4), 914–927.

60. Rolls, R. J., Deane, D. C., Johnson, S. E., Heino, J., Anderson, M. J., & Ellingsen, K. E. (2023). Biotic homogenisation and differentiation as directional change in beta diversity: Synthesising driver–response relationships to develop conceptual models across ecosystems. Biological Reviews, 98(4), 1388–1423. 10.1111/brv.12958

61. Rourke, M. L., Fowler, A. M., Hughes, J. M., Broadhurst, M. K., DiBattista, J. D., Fielder, S., Wilkes Walburn, J., & Furlan, E. M. (2022). Environmental DNA (eDNA) as a tool for assessing fish biomass: A review of approaches and future considerations for resource surveys. Environmental DNA, 4(1), 9–33. 10.1002/edn3.185

62. Schinegger, R., Trautwein, C., Melcher, A., & Schmutz, S. (2012). Multiple human pressures and their spatial patterns in European running waters. Water and Environment Journal, 26(2), 261–273. 10.1111/j.1747-6593.2011.00285.x

63. Schmidt-Kloiber, A., & Hering, D. (2019). Freshwaterecology.info – An Online Database for European Freshwater Organisms, their Biological Traits and Ecological Preferences. Biodiversity Information Science and Standards, 3, e37379. 10.3897/biss.3.37379

64. Schulze, T., Ricking, M., Schröter-Kermani, C., Körner, A., Denner, H.-D., Weinfurtner, K., Winkler, A., & Pekdeger, A. (2007). The German Environmental Specimen Bank. Journal of Soils and Sediments, 7(6), 361–367. 10.1065/jss2007.08.248

65. Schütz, R., Fukaya, K., Beermann, A. J., Díaz, C., Lenz, A.-C., Macher, T.-H., Koschorreck, J., & Leese, F. (2025). Optimizing Environmental DNA Metabarcoding of Archived Suspended Particulate Matter for River Biodiversity Monitoring. Environmental DNA, 7(4), e70130. 10.1002/edn3.70130

66. Schweizer, M., Dieterich, A., Betz, S., Leim, D., Prozmann, V., Jacobs, B., Wick, A., Köhler, H.-R., & Triebskorn, R. (2022). Fish health in the Nidda as an indicator for ecosystem integrity: A case study for Central European small streams in densely populated areas. Environmental Sciences Europe, 34(1), 10. 10.1186/s12302-021-00584-x

67. Seymour, M., Edwards, F. K., Cosby, B. J., Bista, I., Scarlett, P. M., Brailsford, F. L., Glanville, H. C., de Bruyn, M., Carvalho, G. R., & Creer, S. (2021). Environmental DNA provides higher resolution assessment of riverine biodiversity and ecosystem function via spatio-temporal nestedness and turnover partitioning. Communications Biology, 4(1), Article 1. 10.1038/s42003-021-02031-2

68. Sullivan, A. R., Karlsson, E., Svensson, D., Brindefalk, B., Villegas, J. A., Mikko, A., Bellieny, D., Siddique, A. B., Johansson, A.-M., Grahn, H., Sundell, D., Norman, A., Esseen, P.-A., Sjödin, A., Singh, N. J., Brodin, T., Forsman, M., & Stenberg, P. (2025). Airborne eDNA captures three decades of ecosystem biodiversity. Nature Communications, 16(1), 11281. 10.1038/s41467-025-67676-7

69. Taberlet, P., Bonin, A., Zinger, L., & Coissac, E. (2018). Environmental DNA: For Biodiversity Research and Monitoring. Oxford University Press. 10.1093/oso/9780198767220.001.0001

70. Thalinger, B., Rieder, A., Teuffenbach, A., Pütz, Y., Schwerte, T., Wanzenböck, J., & Traugott, M. (2021). The Effect of Activity, Energy Use, and Species Identity on Environmental DNA Shedding of Freshwater Fish. Frontiers in Ecology and Evolution, 0. 10.3389/fevo.2021.623718

71. Theilen, J., Sarrazin, V., Hauten, E., Koll, R., Möllmann, C., Fabrizius, A., & Thiel, R. (2025). Environmental factors shaping fish fauna structure in a temperate mesotidal estuary: Periodic insights from the Elbe estuary across four decades. *Estuarine*, Coastal and Shelf Science, 318, 109208. 10.1016/j.ecss.2025.109208

72. Thomsen, P. F., Kielgast, J. O. S., Iversen, L. L., Wiuf, C., Rasmussen, M., Gilbert, M. T. P., Orlando, L., & Willerslev, E. (2012). Monitoring endangered freshwater biodiversity using environmental DNA. Molecular Ecology, 21(11), 2565–2573.

73. Tickner, D., Opperman, J. J., Abell, R., Acreman, M., Arthington, A. H., Bunn, S. E., Cooke, S. J., Dalton, J., Darwall, W., Edwards, G., Harrison, I., Hughes, K., Jones, T., Leclère, D., Lynch, A. J., Leonard, P., McClain, M. E., Muruven, D., Olden, J. D., … Young, L. (2020). Bending the Curve of Global Freshwater Biodiversity Loss: An Emergency Recovery Plan. BioScience, 70(4), 330–342. 10.1093/biosci/biaa002

74. van der Oost, R., Beyer, J., & Vermeulen, N. P. E. (2003). Fish bioaccumulation and biomarkers in environmental risk assessment: A review. Environmental Toxicology and Pharmacology, 13(2), 57–149. 10.1016/S1382-6689(02)00126-6

75. Vos, M., Hering, D., Gessner, M. O., Leese, F., Schäfer, R. B., Tollrian, R., Boenigk, J., Haase, P., Meckenstock, R., Baikova, D., Bayat, H., Beermann, A., Beisser, D., Beszteri, B., Birk, S., Boden, L., Brauer, V., Brauns, M., Buchner, D., … Sures, B. (2023). The Asymmetric Response Concept explains ecological consequences of multiple stressor exposure and release. Science of The Total Environment, 872, 162196. 10.1016/j.scitotenv.2023.162196

76. Walsh, C. J., Roy, A. H., Feminella, J. W., Cottingham, P. D., Groffman, P. M., & Morgan, R. P. (2005). The urban stream syndrome: Current knowledge and the search for a cure. Journal of the North American Benthological Society, 24(3), 706–723.

77. Wang, B., Jiao, L., Ni, L., Wang, M., & You, P. (2024). Bridging the gap: The integration of eDNA techniques and traditional sampling in fish diversity analysis. Frontiers in Marine Science, 11. 10.3389/fmars.2024.1289589

78. Yates, M. C., Fraser, D. J., & Derry, A. M. (2019). Meta-analysis supports further refinement of eDNA for monitoring aquatic species-specific abundance in nature. Environmental DNA, 1(1), 5–13. 10.1002/edn3.7

79. Zieritz, A., Gallardo, B., Baker, S. J., Britton, J. R., van Valkenburg, J. L. C. H., Verreycken, H., & Aldridge, D. C. (2017). Changes in pathways and vectors of biological invasions in Northwest Europe. Biological Invasions, 19(1), 269–282. 10.1007/s10530-016-1278-z

80. Zizka, V. M. A., Koschorreck, J., Khan, C. C., & Astrin, J. J. (2022). Long-term archival of environmental samples empowers biodiversity monitoring and ecological research. Environmental Sciences Europe, 34(1), 40. 10.1186/s12302-022-00618-y

