## Supplementary material for "Unraveling long-term trends and drivers of fish biodiversity change using environmental DNA metabarcoding of archived samples": Methods

**Supplement 1**

**S1.1 Methods: Species richness and group trends**

A map of sampling locations was generated using high-resolution administrative boundaries from Natural Earth's 1:10,000,000 scale dataset (Massicotte et al., 2025) and major river networks obtained from Natural Earth for the Rhine, Elbe, and Danube rivers, and from OpenStreetMap (OSM) for the Saar, Saale, Mulde, and Mosel rivers (Padgham et al., 2017). All spatial data were processed in the WGS84 coordinate reference system (EPSG:4326) using the sf package (Pebesma, 2018) and visualized with ggplot2 (Wickham, 2016) in R Studio (RStudio 2025.05.1+513).

Species richness of each site was determined from presence/absence data (reads > 0) and temporal trends in species richness were evaluated using complementary statistical approaches. Mann-Kendall test was used to provide Kendall's tau (τ) for detecting monotonic trends in time-series data (Kendall, 1975; Mann, 1945). To account for non-linear temporal patterns, Generalized Additive Models (GAMs) were fitted using the mgcv package (Wood, 2017) with Poisson regression. As smooth terms we employed cubic regression splines with shrinkage (bs = "cs") and Restricted Maximum Likelihood (REML) estimation. Effective degrees of freedom (edf) near 1 indicate approximately linear relationships, while higher values suggest non-linear temporal patterns. Statistical significance was set at α = 0.05 for all tests. Data manipulation and analysis were performed using dplyr (Wickham et al., 2023), tidyr (Wickham, Vaughan, et al., 2025), readxl (Wickham, Bryan, et al., 2025), openxlsx (Schauberger et al., 2025), Kendall (McLeod, 2022), and trend (Pohlert, 2023) packages in R.

Species were then classified into three functional groups for community analysis. First, species were categorized by conservation status (common native species, red list species, invasive species) based on the German Red List (Freyhof et al., 2023). Second, species were classified by water quality tolerance (sensitive species, tolerant species) using indicator status from freshwaterecology.info (Schmidt-Kloiber & Hering, 2019). Third, species were assigned to migratory behavior categories (migratory, non-migratory) also from freshwaterecology.info (Schmidt-Kloiber & Hering, 2019). We then calculated three community metrics: species richness (absolute count of species), proportional species richness (group richness divided by total site richness in a year), and proportional read abundance (group reads divided by total site reads per year). Temporal trends were assessed using two approaches for each metric: linear regression with year as the predictor, providing slope estimates representing annual rate of change, and Kendall's τ rank correlation coefficient to evaluate monotonic trends without assuming linearity (Kendall, 1975). Kendall's τ was calculated using the cor.test function with method = "kendall" and exact = FALSE for asymptotic p-value approximation. Statistical significance was assessed at α = 0.05 for both parametric (linear regression) and non-parametric (Kendall's τ) tests. All analyses were conducted in R Studio using tidyverse (Wickham et al., 2019), readxl (Wickham, Bryan, et al., 2025), and writexl (Ooms et al., 2025) packages.

**S1.2 Methods: Beta Diversity analysis**

The community composition was analyzed using both quantitative (relative read abundance) and qualitative (presence/absence) approaches. Beta diversity was quantified using Bray-Curtis dissimilarity for abundance comparisons and Jaccard dissimilarity for presence/absence data (Bray & Curtis, 1957; Jaccard, 1912). Dissimilarity indices were calculated using the vegan package (Oksanen et al., 2025). To evaluate temporal turnover and spatial homogenization patterns, we calculated within-site temporal dissimilarity, measuring compositional change at individual sites across time lags, and between-site spatial dissimilarity, measuring changes in spatial heterogeneity across the sites over time. Within-site dissimilarity was calculated for all pairwise comparisons between years at the same site. Between-site dissimilarity was calculated as the change in mean dissimilarity between sites at different time lags. Beta diversity was further investigated by calculating turnover and nestedness using the betapart package (Baselga & Orme, 2012). Temporal trends in dissimilarity metrics were assessed using Kendall's τ rank correlation to evaluate monotonic changes over time lags.

The community diversity was further quantified using Hill numbers q = 0, 1, and 2, which provide a unified framework for measuring species diversity (Chao et al., 2014; Hill, 1973). Hill number q = 0 equals species richness, q = 1 equals the exponential of Shannon entropy and represents the effective number of common species, and q = 2 equals the inverse of Simpson concentration and represents the effective number of dominant species. Diversity stability was assessed for each site using the coefficient of variation (CV) of Hill number q = 1 across all years. Sites were classified into four stability categories based on median diversity (high/low) and CV thresholds (stable: CV ≤ 0.15; unstable: CV > 0.15), following established frameworks for ecosystem stability assessment (Donohue et al., 2016). Temporal trends in Hill numbers at each site were evaluated using Kendall's τ correlation and linear regression.

Species-level temporal trends in relative abundance were analyzed across all sites using linear regression. Kendall's τ correlation was calculated to assess monotonic trends, with statistical significance evaluated at α = 0.05. All analyses were conducted in R Studio using tidyverse (Wickham et al., 2019), vegan (Oksanen et al., 2025), betapart (Baselga & Orme, 2012), and patchwork (Pedersen, 2025) packages. Visualizations were done using ggplot2 (Wickham, 2016) and ggrepel (Slowikowski et al., 2024).

**S1.3 Methods: Correlation between relative reads and site occupancy**

To assess the relationship between trends in spatial occurrence and read abundance trends for each species, we calculated site occupancy, defined as the proportion of sampled sites where the species was detected (presence/absence) and relative read abundance, calculated as the proportion of reads represented by each species. Species-specific temporal trends were assessed using Mann-Kendall's tau (τ) statistic (Kendall, 1975; Mann, 1945). The tau values were calculated independently for both occupancy and relative abundance using the Kendall package (McLeod, 2022). Statistical significance was evaluated at α = 0.05, and species were classified into four categories based on their trend significance. Significant trends in both metrics, significant trends in occupancy only, significant trends in relative abundance only, or no significant trends. The correlation between occupancy trends and relative abundance trends was assessed using Spearman's rank correlation. The relationship was visualized using a quadrant scatterplot with occupancy τ values on the x-axis and relative abundance τ values on the y-axis, divided into four quadrants representing the different scenarios. Quadrant I (expanding range with increasing abundance), Quadrant II (decreasing range with increasing abundance), Quadrant III (decreasing range with decreasing abundance), and Quadrant IV (expanding range with decreasing abundance). A linear regression line with 95% confidence interval was fitted to quantify the overall relationship. All analyses were conducted in R Studio using readxl (Wickham, Bryan, et al., 2025), dplyr and tidyr (Wickham et al., 2023; Wickham, Vaughan, et al., 2025), Kendall (McLeod, 2022), openxlsx (Schauberger et al., 2025), ggplot2 (Wickham, 2016), ggrepel (Slowikowski et al., 2024), and stringr (Wickham, Software, et al., 2025).

**S1.4 Methods: eDNA trend estimates**

Temporal trends in fish species relative abundance were analyzed using Bayesian hierarchical models with the brms package (Bürkner, 2017) in R Studio, similar to the analysis of Friedrichs-Manthey et al., 2024 with dataset-specific differences. Multiple OTUs assigned to the same species were aggregated by summing read counts within each site-year combination. Relative read abundance was calculated by dividing each species' read count by the total read count per sample, then logit-transformed to normalize distributions and stabilize variance. For each species with observations from at least two different years, we fitted a mixed-effects model with site as a random intercept (or random slope when data permitted). Models used Gaussian error distribution on logit-transformed responses. Non-detections were included as zeros with pseudocount adjustments (0.00001 for zeros, 0.99999 for ones) before transformation. Models were then fitted using Markov Chain Monte Carlo (MCMC) sampling with four chains of 2,000-4,000 iterations (including 500-1,000 warmup samples), yielding 6,000-12,000 posterior samples. Convergence was assessed using R-hat ≤ 1.1 (Gelman & Rubin, 1992) and adapt_delta was set to 0.95-0.99. Species trends were classified based on 95% credible intervals of the year coefficient. Significantly increasing (CI entirely > 0), significantly decreasing (CI entirely < 0), or stable (CI overlaps 0). All analyses used tidyverse (Wickham et al., 2019), tidybayes (Kay & Mastny, 2024), and RStan backend (Guo et al., 2025).

**1.5 Methods: Comparison to long-term trend data from regulatory monitoring**

The trend estimates from both methods were expressed as the annual rates of change on the logit scale. Agreement between methods was assessed using multiple complementary approaches. Directional concordance (percentage of species showing the same trend direction), Cohen's kappa (κ) to quantify directional agreement beyond chance (Landis & Koch, 1977), Spearman's rank correlation between trend magnitudes, Wilcoxon rank test for systematic bias, and Bland-Altman analysis to assess mean bias and 95% limits of agreement (Bland & Altman, 1986). Results were visualized using dumbbell charts colored by agreement status and Bland-Altman plots to identify systematic biases. All analyses were conducted in R Studio using tidyverse (Wickham et al., 2019), psych (Revelle, 2025), and patchwork (Pedersen, 2025).

**S1.6. Methods: Physicochemical correlations**

Chemistry data were obtained from the same SPM samples used for metabarcoding (N=66) and complemented with additional water parameters (N=17) collected from the same long-term monitoring sites. The data was collected from public databases (umwelprobenbank.de, undine.bafg.de, gkd.bayern.de, fgg-rhein.bafg.de, elbe-datenportal.de, iksr.bafg.de, umweltdaten.lubw.baden-wuerttemberg.de). In total, 83 environmental parameters were analyzed, including multiple stressor categories. Per- and polyfluoroalkyl substances (PFAS, n=19), heavy metals and trace elements (n=13) including mercury, lead, and arsenic, polycyclic aromatic hydrocarbons (PAHs, n=11), pesticides and persistent organic pollutants (POPs, n=13), pharmaceuticals (n=3), nutrients (n=7) including nitrogen and phosphorus, ions and water chemistry (n=6) including conductivity and mineral composition, physical parameters (n=7) including sediment grain size, temperature, oxygen, and discharge, organic and inorganic carbon (n=3), and stable isotopes (n=2). Annual mean values were calculated for each site and year to match the spatio- temporal resolution of the eDNA data.

The relationship between species richness and individual environmental parameters was assessed using Spearman's rank correlation coefficient calculated using Fisher's z-transformation. Only chemical parameters with at least 30 observations were included. Correlations were considered significant at α= 0.05. The species-level relationships between relative abundance and chemical parameters were assessed similarly, requiring a minimum of 10 observations per species-chemical combination.

To assess multivariate relationships between community composition and chemical gradients, we first evaluated gradient length using Detrended Correspondence Analysis (DCA). As gradient length was below 3 standard deviations, Redundancy Analysis (RDA) was used. Community data were Hellinger-transformed prior to RDA to account for the lack of absences (zeros) in species data (Legendre & Gallagher, 2001). Multicollinearity among chemical predictors was assessed using Variance Inflation Factors (VIF). Forward selection using the ordiR2step() function identified the most parsimonious set of chemical predictors that significantly explained compositional variation, using adjusted R² as the optimization criterion and permutation tests (n = 999) for variable significance. The significance of the overall ordination model and individual axes was assessed using a PERMANOVA (n=999). To distinguish chemical effects from spatial and temporal confounding, we performed partial RDA (pRDA) using site identity and year as covariates. For this we categorized chemicals per site (tests temporal chemical effects within sites), chemicals per year (tests spatial chemical effects among sites), and chemicals on both site and year (tests chemical effects independent of spatiotemporal structure).

To evaluate the predictive power of chemical parameters, Random Forest (RF) models were fitted using leave-one-site-out cross-validation (LOSOCV) to ensure independence between training and test sets. For richness prediction, a single RF model with 500 trees was trained to predict species richness from standardized chemical predictors. Model performance was evaluated using R², root mean squared error (RMSE) and mean absolute error (MAE). Variable importance was quantified using permutation importance, which measures the increase in prediction error when each predictor is randomly shuffled. Statistical significance of RF models was assessed using permutation tests (n=999). The entire response variable was permuted, and models were refitted to generate a null distribution of R² values. Observed R² values were compared to null distributions to calculate p-values. All analyses were conducted in R using vegan (Oksanen et al., 2025) for ordination, ranger (Wright & Ziegler, 2017) for Random Forest, and tidyverse (Wickham et al., 2019) for data manipulation and visualization.

**References**

Baselga, A., & Orme, C. D. L. (2012). betapart: An R package for the study of beta diversity. *Methods in Ecology and Evolution*, *3*(5), 808–812. https://doi.org/10.1111/j.2041-210X.2012.00224.x

Bland, J. M., & Altman, D. (1986). STATISTICAL METHODS FOR ASSESSING AGREEMENT BETWEEN TWO METHODS OF CLINICAL MEASUREMENT. *The Lancet*, *327*(8476), 307–310. https://doi.org/10.1016/S0140-6736(86)90837-8

Bray, J. R., & Curtis, J. T. (1957). An Ordination of the Upland Forest Communities of Southern Wisconsin. *Ecological Monographs*, *27*(4), 325–349. https://doi.org/10.2307/1942268

Bürkner, P.-C. (2017). brms: An R Package for Bayesian Multilevel Models Using Stan. *Journal of Statistical Software*, *80*, 1–28. https://doi.org/10.18637/jss.v080.i01

Chao, A., Chiu, C.-H., & Jost, L. (2014). Unifying Species Diversity, Phylogenetic Diversity, Functional Diversity, and Related Similarity and Differentiation Measures Through Hill Numbers. *Annual Review of Ecology, Evolution, and Systematics*, *45*(Volume 45, 2014), 297–324. https://doi.org/10.1146/annurev-ecolsys-120213-091540

Donohue, I., Hillebrand, H., Montoya, J. M., Petchey, O. L., Pimm, S. L., Fowler, M. S., Healy, K., Jackson, A. L., Lurgi, M., McClean, D., O’Connor, N. E., O’Gorman, E. J., & Yang, Q. (2016). Navigating the complexity of ecological stability. *Ecology Letters*, *19*(9), 1172–1185. https://doi.org/10.1111/ele.12648

Freyhof, J.; Bowler, D.; Broghammer, T.; Friedrichs-Manthey, M.; Heinze, S. & Wolter, C. (2023). *Rote Liste und Gesamtartenliste der sich im Süßwasser reproduzierenden Fische und Neunaugen (Pisces et Cyclostomata) Deutschlands*. Landwirtschaftsverlag GmbH. https://doi.org/10.19213/972176

Friedrichs-Manthey, M., Bowler, D. E., & Freyhof, J. (2024). Freshwater fish in mid and northern German rivers – Long-term trends and associated species traits. *Science of The Total Environment*, *957*, 177759. https://doi.org/10.1016/j.scitotenv.2024.177759

Gelman, A., & Rubin, D. B. (1992). Inference from Iterative Simulation Using Multiple Sequences. *Statistical Science*, *7*(4), 457–472. https://doi.org/10.1214/ss/1177011136

Guo, J., Gabry, J., Goodrich, B., Johnson, A., Weber, S., Badr, H. S., Lee, D., Sakrejda, K., Martin, M., University, T. of C., Sklyar (R/cxxfunplus.R), O., Team (R/pairs.R, T. R. C., R/dynGet.R), Oehlschlaegel-Akiyoshi (R/pairs.R), J., Maddock (gamma.hpp), J., Bristow (gamma.hpp), P., Agrawal (gamma.hpp), N., Kormanyos (gamma.hpp), C., & Steve, B. (2025). *rstan: R Interface to Stan* (Version 2.32.7) [Computer software]. https://cran.r-project.org/web/packages/rstan/index.html

Hill, M. O. (1973). Diversity and Evenness: A Unifying Notation and Its Consequences. *Ecology*, *54*(2), 427–432. https://doi.org/10.2307/1934352

Jaccard, P. (1912). The Distribution of the Flora in the Alpine Zone. *New Phytologist*, *11*(2), 37–50. https://doi.org/10.1111/j.1469-8137.1912.tb05611.x

Kay, M., & Mastny, T. (2024). *tidybayes: Tidy Data and “Geoms” for Bayesian Models* (Version 3.0.7) [Computer software]. https://cran.r-project.org/web/packages/tidybayes/index.html

Kendall, M. G. (1975). *Rank correlation methods* (4th ed., 2d impression). Griffin.

Landis, J. R., & Koch, G. G. (1977). The measurement of observer agreement for categorical data. *Biometrics*, *33*(1), 159–174.

Legendre, P., & Gallagher, E. D. (2001). Ecologically meaningful transformations for ordination of species data. *Oecologia*, *129*(2), 271–280. https://doi.org/10.1007/s004420100716

Mann, H. B. (1945). Nonparametric Tests Against Trend. *Econometrica*, *13*(3), 245–259. https://doi.org/10.2307/1907187

Massicotte, P., South, A., & Hufkens, K. (2025). *rnaturalearth: World Map Data from Natural Earth* (Version 1.1.0) [Computer software]. https://cran.r-project.org/web/packages/rnaturalearth/index.html

McLeod, A. I. (2022). *Kendall: Kendall Rank Correlation and Mann-Kendall Trend Test* (Version 2.2.1) [Computer software]. https://cran.r-project.org/web/packages/Kendall/index.html

Oksanen, J., Simpson, G. L., Blanchet, F. G., Kindt, R., Legendre, P., Minchin, P. R., O’Hara, R. B., Solymos, P., Stevens, M. H. H., Szoecs, E., Wagner, H., Barbour, M., Bedward, M., Bolker, B., Borcard, D., Borman, T., Carvalho, G., Chirico, M., Caceres, M. D., … Weedon, J. (2025). *vegan: Community Ecology Package* (Version 2.7-2) [Computer software]. https://cran.r-project.org/web/packages/vegan/index.html

Ooms [aut, J., cre, & details, J. M. (Author of libxlsxwriter (see A. and C. files for details)) writexl author. (2025). *writexl: Export Data Frames to Excel “xlsx” Format* (Version 1.5.4) [Computer software]. https://cran.r-project.org/web/packages/writexl/index.html

Padgham, M., Lovelace, R., Salmon, M., & Rudis, B. (2017). Osmdata. *Journal of Open Source Software*, *2*(14), 305. https://doi.org/10.21105/joss.00305

Pebesma, E. (2018). Simple Features for R: Standardized Support for Spatial Vector Data. *The R Journal*, *10*(1), 439–446. https://doi.org/10.32614/RJ-2018-009

Pedersen, T. L. (2025). *patchwork: The Composer of Plots* (Version 1.3.2) [Computer software]. https://cran.r-project.org/web/packages/patchwork/index.html

Pohlert, T. (2023). *trend: Non-Parametric Trend Tests and Change-Point Detection* (Version 1.1.6) [Computer software]. https://cran.r-project.org/web/packages/trend/index.html

Revelle, W. (2025). *psych: Procedures for Psychological, Psychometric, and Personality Research* (Version 2.5.6) [Computer software]. https://cran.r-project.org/web/packages/psych/index.html

Schauberger, P., Walker, A., Braglia, L., Sturm, J., Garbuszus, J. M., Barbone, J. M., Zimmermann, D., & Kainhofer, R. (2025). *openxlsx: Read, Write and Edit xlsx Files* (Version 4.2.8.1) [Computer software]. https://cran.r-project.org/web/packages/openxlsx/index.html

Schmidt-Kloiber, A., & Hering, D. (2019). Freshwaterecology.info – An Online Database for European Freshwater Organisms, their Biological Traits and Ecological Preferences. *Biodiversity Information Science and Standards*, *3*, e37379. https://doi.org/10.3897/biss.3.37379

Slowikowski, K., Schep, A., Hughes, S., Dang, T. K., Lukauskas, S., Irisson, J.-O., Kamvar, Z. N., Ryan, T., Christophe, D., Hiroaki, Y., Gramme, P., Abdol, A. M., Barrett, M., Cannoodt, R., Krassowski, M., Chirico, M., Aphalo, P., & Barton, F. (2024). *ggrepel: Automatically Position Non-Overlapping Text Labels with “ggplot2”* (Version 0.9.6) [Computer software]. https://cran.r-project.org/web/packages/ggrepel/index.html

Wickham, H. (2016). *Ggplot2*. Springer International Publishing. https://doi.org/10.1007/978-3-319-24277-4

Wickham, H., Averick, M., Bryan, J., Chang, W., McGowan, L. D., François, R., Grolemund, G., Hayes, A., Henry, L., Hester, J., Kuhn, M., Pedersen, T. L., Miller, E., Bache, S. M., Müller, K., Ooms, J., Robinson, D., Seidel, D. P., Spinu, V., … Yutani, H. (2019). Welcome to the Tidyverse. *Journal of Open Source Software*, *4*(43), 1686. https://doi.org/10.21105/joss.01686

Wickham, H., Bryan, J., Posit, attribution), P. (Copyright holder of all R. code and all C. code without explicit copyright, code), M. K. (Author of included R., code), K. V. (Author of included libxls, code), C. L. (Author of included libxls, code), B. C. (Author of included libxls, code), D. H. (Author of included libxls, & code), E. M. (Author of included libxls. (2025). *readxl: Read Excel Files* (Version 1.4.5) [Computer software]. https://cran.r-project.org/web/packages/readxl/index.html

Wickham, H., François, R., Henry, L., Müller, K., Vaughan, D., Software, P., & PBC. (2023). *dplyr: A Grammar of Data Manipulation* (Version 1.1.4) [Computer software]. https://cran.r-project.org/web/packages/dplyr/index.html

Wickham, H., Software, P., & PBC. (2025). *stringr: Simple, Consistent Wrappers for Common String Operations* (Version 1.6.0) [Computer software]. https://cran.r-project.org/web/packages/stringr/index.html

Wickham, H., Vaughan, D., Girlich, M., Ushey, K., Software, P., & PBC. (2025). *tidyr: Tidy Messy Data* (Version 1.3.2) [Computer software]. https://cran.r-project.org/web/packages/tidyr/index.html

Wood, S. N. (2017). *Generalized Additive Models: An Introduction with R, Second Edition* (2nd ed.). Chapman and Hall/CRC. https://doi.org/10.1201/9781315370279

Wright, M. N., & Ziegler, A. (2017). ranger: A Fast Implementation of Random Forests for High Dimensional Data in C++ and R. *Journal of Statistical Software*, *77*, 1–17. https://doi.org/10.18637/jss.v077.i01
